# Mitofusin agonists improve the long-term repopulating activity of human cord blood HSCs after *ex vivo* culture

**DOI:** 10.1101/2025.10.27.684807

**Authors:** Alyssa Biondo, Gabriela Candelaria, Daniel K. Jin, Emily Lin, Daniel J. McLaughlin, Jessica Freedman, Zhaolun Liang, Katherine J. Leong, Ya-Ju Chang, Zhiran Geng, Nazia Nayeem, Zachary Leontiou, Emma Laquinta, Christopher D. Hillyer, Hans-Willem Snoeck, Larry L. Luchsinger

## Abstract

Umbilical cord blood (CBU) is a valuable source of hematopoietic stem cells (HSCs) due to lower incidence of graft-versus-host disease. However, limited HSC counts restrict use in adults, necessitating improved methods for *ex vivo* expansion and improved HSC function. Mitofusin 2 (MFN2), a mitochondrial membrane fusion protein, is necessary for preserving HSC function and mitofusin agonists have been characterized. We report ex vivo culture of CBU HSCs with mitofusin agonists (MAs) enhances long-term repopulating activity by over five-fold in both primary and secondary transplantation assays without changes of total nucleated cells or phenotypic HSCs. Mechanistically, MA-treated HSCs show suppressed protein synthesis, increased autophagic flux, and elevated lysosomal mass. Transcriptomic analysis implicates downregulation of MTOR signaling, and immunoprecipitation studies suggest MFN2 interacts with MTOR. These data support a model in which fusion-competent MFN2 sequesters MTOR, promoting a catabolic state that preserves HSC potency. Our findings suggest a novel MFN2–MTOR regulatory axis that enhances the function of human HSCs cultured *ex vivo* for potential therapeutic application.

## Introduction

Human cord blood (CB) and bone marrow (BM) contain hematopoietic stem cells (HSCs) that are capable of long-term reconstitution of peripheral blood in patients receiving myeloablative therapy.^1^ CB units have clinical advantages in their ease of availability and high fraction of naïve lymphocytes which produce less activated T lymphocytes and reduce the risk of graft-versus-host-disease (GVHD), a major hurdle in successful long-term HSC transplantation.^2–4^ Expansion of CBU cells *ex vivo* for periods sufficient to increase nucleated cell counts to clinically useful levels remains a significant challenge due to reduced long-term HSC function. While several small molecules such as UM171, SR1 and nicotinamide (NAM) analogs have been demonstrated to increase the total nuclear cells and functional HSCs after *ex vivo* culture, expanded CBUs have not yet been adopted clinically.^5–8^ Thus, elucidating novel mechanisms of HSC maintenance and self-renewal is necessary to develop novel methods to sustain long-term engraftment towards patient-specific HSC expansion and improved transplantation outcomes.

The mitochondrial fusion protein Mitofusin 2 (MFN2) has been shown to be required for HSC function in mice.^9^ Transactivation of MFN2 via the transcription factor Prdm16 induces high mitochondrial fusion in the HSC compartment and conditional deletion of MFN2 in the hematopoietic system reduces HSC potency and skews lineage output towards a myeloid biased phenotype in transplantation assays. Recently, mitofusin agonists (MAs) have been developed to induce a fusion competent conformation of mitofusins independent of endogenous GTP hydrolysis activity and increase MFN2 fusion activity.^10^

In this study, we identify mitochondrial agonists (MAs) as effective agents for improving long-term engraftment and potency of hematopoietic stem cell (HSC) after *ex vivo* culture. Our research reveals a mechanism by which MAs attenuate mammalian target of rapamycin (MTOR) signaling, resulting in reduced protein synthesis and elevated autophagic flux in HSC during *ex vivo* culture: hallmarks of potent HSCs.^11,12^ Our findings present a novel strategy to mitigate exhaustion of HSCs during *ex vivo* culture, potentially improving clinical utility and enhancing hematopoietic stem cell transplantation (HSCT) outcomes for patients.

## Methods

### Animals

NOD.Cg-Prkdc^scid^ Il2rg^tm1Wjl^/SzJ were purchased from The Jackson Laboratory. Animals were housed in a pathogen-free facility. Experiments and animal care were performed in accordance with the New York Blood Center Institutional Animal Care and Use Committee. The mice used for transplant were between 8 – 12 weeks of age. Randomly chosen littermates of both sexes were used for transplantation.

### Cryopreserved Cord Blood Unit HSC Isolation

De-identified human cord blood units (CBU) were obtained from the National Cord Blood Program of the New York Blood Center. CBUs were in LN2 vapor freezers until required and mononuclear cells were used as described (Supplemental Materials).

### Hematopoietic Stem Cell Transplantation

After 7 days of culture, NSG mice were sub-lethally irradiated NSG with 150cGy (120cGy/min) using an XRad320 source (Precision X-Ray). Mice were transplanted by tail vein injection with 25% of HSC cultures in a volume of 200μL into each recipient. Human peripheral blood chimerism were monitored by flow cytometry at 8, 15, 25 and 30 weeks after transplantation. At 30 weeks mice were sacrificed, and BM cells were collected by the crushing method above. Total BM cells were analyzed by flow cytometry. For secondary transplantation, 1x10^6^ BM cells from 30-week primary recipients were transplanted into secondary sub-lethally irradiated NSG recipients and human peripheral blood chimerism were monitored by flow cytometry at the indicated time points. For limiting dilution assays, NSG recipients were transplanted with high dose (4,000) or low dose (2,000) total cultured cells after 7-day cultures. Peripheral blood was analyzed for human chimerism 15 weeks post-transplant. HSC frequencies were calculated using the ELDA software based on the number of non-repopulated mice defined as ≤0.1% donor contribution.^13^

### RNA-seq and Gene Set Enrichment Analysis from HSCs

HSC cultures were resorted for phenotypic CD90+ HSCs and lysed in Trizol LS overnight at -80C and mRNA was extracted according to manufacturer’s protocol. Amplification of cDNA was carried out using Superscript III reverse transcriptase and resuspended in 12uL of diH2O. Sequencing libraries were prepared using an SMartR 2 RNA-seq system V2 kit. Paired-end 150 bp sequencing was performed on an Illumina HiSeq 4000. Alignment to the human genome (hg37) was performed BasePair (https://basepairtech.com). Expression counts were normalized to FPKM.

### Statistics

Statistical analyses were performed using Prism 10 software. The unpaired student’s t-test was used for statistical analysis between 2 groups. For parametric analyses, the one-way ANOVA with Dunnett’s test was used for comparison between ≥3 groups. For non-parametric analyses, the Kruskal-Wallace test was used for multiple comparisons between more than 2 groups. The Brown-Forsythe and Welsch ANOVA test with the Games-Howell multiple comparisons post-hoc test was used for data with unequal standard deviations or sample sizes. Results are expressed as mean ± SEM. Prism 10 (GraphPad) was used for calculations. A *p* value of less than 0.05 was defined as statistically significant. Sample size represents biological replicates. Cochran’s Q test was used for exclusion of outliers.

## Results

### Elevated MFN2 expression and mitochondrial fusion characterize CBU HSCs

Cord blood units (CBU) were used as a clinically relevant HSC source because of their potency, availability, and underutilization in transplantation.^14^ We began by examining MFN2 expression in CBU progenitor cell compartments. Randomized cryopreserved CBUs from the National Cord Blood Program were used to isolate HSCs (i.e. CD90^+^ HSCs, Lin^-^CD34^+^CD38^-^CD45RA^-^CD90^+^), multipotent progenitors (MPPs, Lin^-^CD34^+^CD38^-^CD45RA^-^CD90^-^), lymphoid-primed multi-potent progenitors (LMPPs, Lin^-^CD34^+^CD38^-^CD45RA^+^CD90^-^), common myeloid progenitors (CMPs, Lin^-^CD34^+^CD38^+^IL3R^+^), and lineage positive cells (Lin^+^) cells for *ex vivo* analyses (**Supplementary Figure S1A**). Quantitative RT-PCR analysis demonstrated enriched *MFN2* expression in HSCs with reduced expression in progenitor and Lin^+^ populations, mirroring expression of inner mitochondrial fusion protein *OPA1* as well as *PRDM16*, a known transactivator of *Mfn2* in murine hematopoiesis. (**Figure 1A**). Notably, expression of the fusion homologue *MFN1* was not significantly altered between progenitor compartments while specific upregulation of mitochondria fission genes *DRP1* and *FIS1* was observed in MPPs but not LMPPs (**Figure 1A, Supplementary Figure S1B**). Intriguingly, further examination of the Lin^+^ population showed ∼89% higher expression of *MFN2* in CD33^+^ myeloid cells compared to HSCs while CD3^+^ T-cells and CD19^+^ B-cells showed significantly lower expression (**Supplementary Fig. S1B**), supporting a previously identified role for MFN2 in innate immune cell function.^15^ These data suggest altered mitochondrial dynamics may describe functional differences of hematopoietic cell populations.^16^

**Figure 1:**
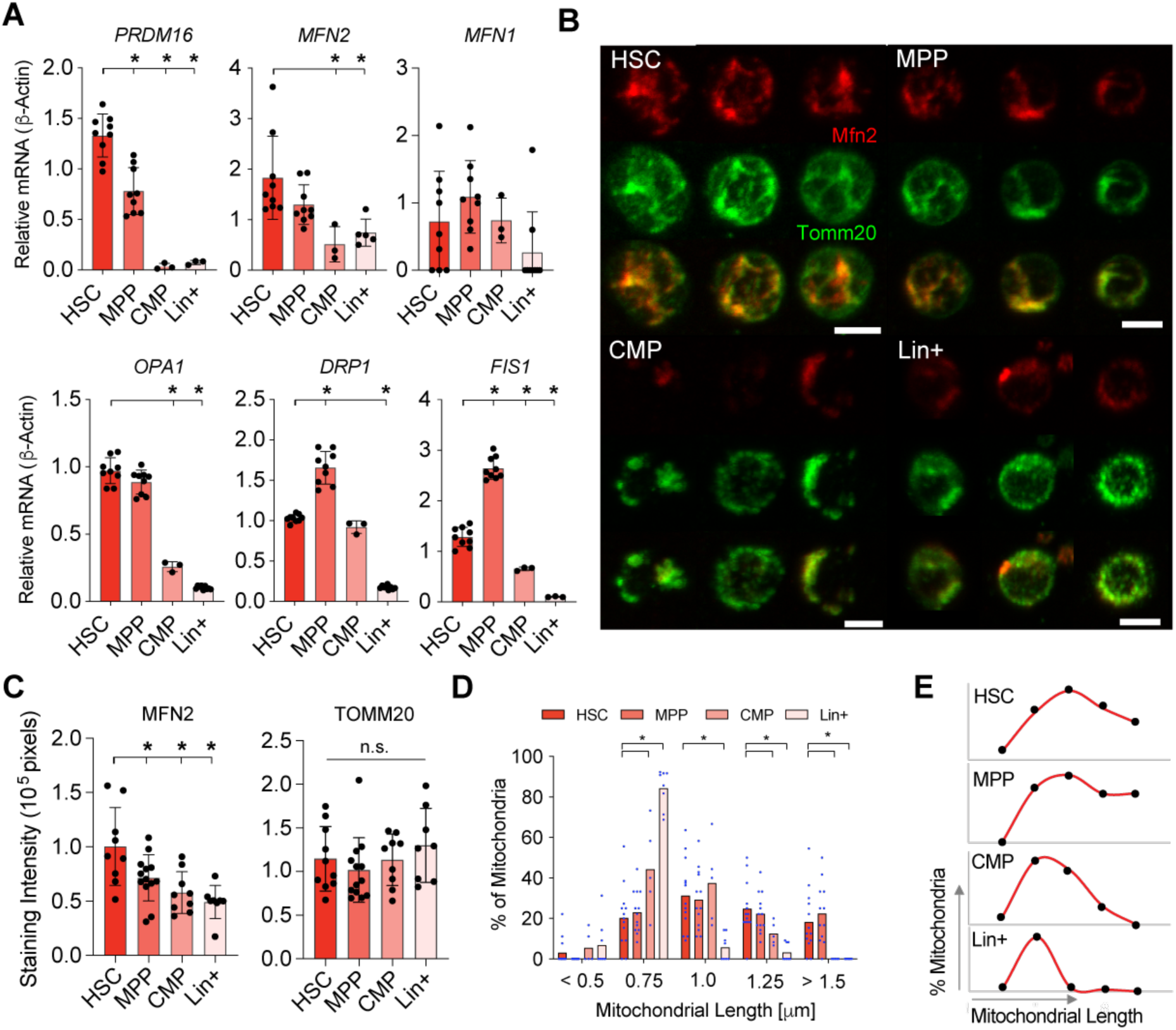
MFN2 is highly expressed in CBU HSCs. **A**, Relative mRNA expression via qRT-PCR of PRDM16, MFN2, MFN1, OPA1, DRP1, and FIS1 gene expression in phenotypic CBU cell populations; n = 3-9 biological replicates; *p < 0.05, Brown-Forsythe and Welch’s ANOVA test with the Games-Howell multiple comparisons post-hoc test for unequal group variances and sample sizes. **B**, Immunofluorescent (IF) antibody staining for MFN2 and TOMM20 in HSC, CMP, MPP, and Lin^+^ cells; scale bar is 5µm, 3 replicate micrographs are shown. **C**, Quantification of IF staining for MFN2 and TOMM20; n = 8-13 fields of cells from three biological replicates; *p < 0.05, one-way ANOVA with Dunnett’s post-hoc test. **D**, Frequency of mitochondrial lengths in isolated CBU cell populations stained with Mitotracker Red (MTR); mean of ≥15 fields of cells and ≥90 mitochondria from ≥8 biological replicates, two-way ANOVA with Tukey’s multiple comparisons test. **E**, Mitochondrial length profiles in CBU HSC, MPP, CMP, and Lin^+^ cells.

Immunofluorescence microscopy confirmed significantly higher MFN2 expression in HSCs than in CBU progenitor and Lin^+^ compartments, with strong colocalization with the mitochondrial marker TOMM20. (**Figure 1B, 1C**). Freshly sorted CBU populations were stained with MitoTracker Red (MTR) and analyzed by super-resolution confocal microscopy (SRCM) to quantify mitochondrial morphology. HSCs and MPPs exhibited longer mitochondria and a higher fusion profile than more differentiated CBU populations (**Figures 1D–E**). To assess MFN2-associated mitochondria–ER contacts, sorted CBU cells were stained for TOMM20 and the ER marker sigma-1 receptor (S1R), as MFN2 mediates mitochondria–ER tethering.^17,18^ SRCM analysis showed significantly greater colocalization of TOMM20 and S1R in HSCs than in CBU progenitor cells. (**Supplementary Figure S1C, S1D**). Taken together, these data suggest elevated MFN2 expression and mitochondrial fusion activity is a feature of human CBU HSCs.

### Pharmacological mitofusin agonists maintain functionally potent HSCs during *ex vivo* culture

MFN2 contains an N-terminal GTPase domain and two C-terminal heptad repeat domains flanking a double-pass transmembrane domain.^19^ Conformational switching between closed, fusion-inhibited and open, fusion-competent states regulates MFN2-mediated mitochondrial tethering and fusion, while mitofusin agonists allosterically promote the fusion-competent conformation independently of GTPase activity (**Figure 2A**).^10,20^ Rocha et al. described both small molecule and peptide MAs that catalyze the MFN2 open conformation of either PINK1 kinase phosphorylated (Cmp A, Pep D) or unphosphorylated (Cmp B, Pep S) MFN2 proteins (**Supplementary Figure S2A**). PINK1 phosphorylation of MFN2 initiates PARKIN-mediated mitophagy and can activate MFN2 fusion to autophagosomes in some tissues. As PINK1 activity is debated in HSC biology,^21,22^ we chose to investigate the effect of both classes of MAs on HSC *ex vivo* culture studies.

**Figure 2:**
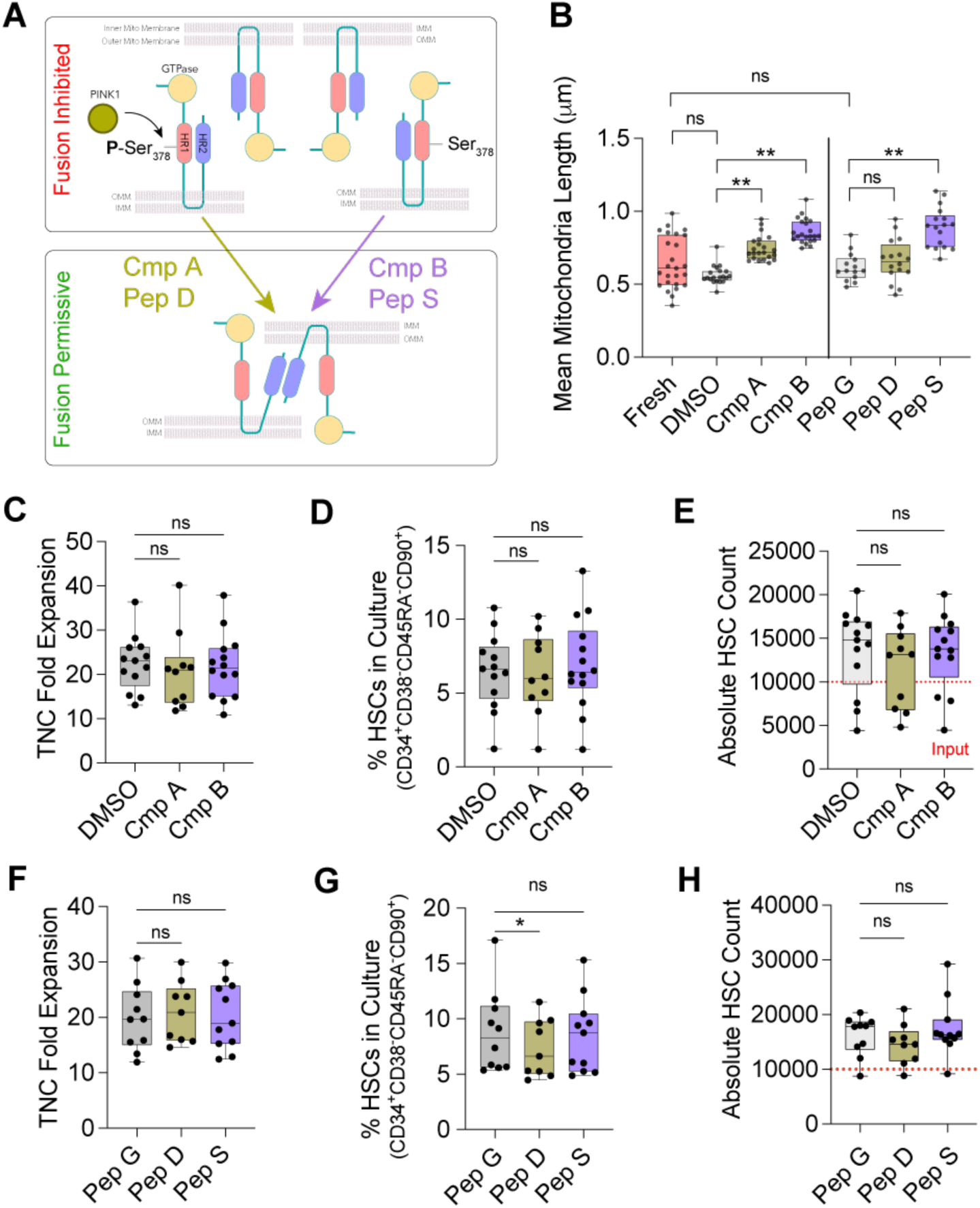
MAs do not affect the phenotypic yield of HSCs following *ex vivo* culture. **A**, Depiction of the fusion inhibited (top) and fusion permissive (bottom) conformation of MFN2 protein. Compound A, Peptide D induce an open MFN2 conformation in the presence of PINK1 phosphorylation of S378, while Compound B and Peptide S induce an open conformation of unphosphorylated MFN2. **B**, Quantification of mitochondria lengths (µm) in freshly isolated or resorted CD90^+^ HSCs cultured for 7 days with DMSO, 5nM Cmp A, 5nM Cmp B, or 1μM Peptide G, 1μM Peptide D, or 1μM Peptide S stained with MTR; n = 15-20 individual cells from three biological replicates, *p<0.05 and **p<0.01, one-way ANOVA with Dunnett’s post-hoc test. **C**, Total nucleated cells (TNC) fold expansion of CD90+ HSC cultures for 7 days with DMSO, 5nM Cmp A or 5nM Cmp B; n = 11-13 independent cultures, *p = 0.05, one-way ANOVA with Dunnett’s post-hoc test. **D**, Percent CD90+ HSCs in culture after 7 days with DMSO, 5nM Cmp A or 5nM Cmp B; n ≥ 11-13 independent cultures, *p = 0.05, one-way ANOVA with Dunnett’s post-hoc test. **E**, Absolute CD90+ HSC counts in cultures after 7 days with DMSO, 5nM Cmp A or 5nM Cmp B; n ≥ 11-13 independent cultures, *p = 0.05, one-way ANOVA with Dunnett’s post-hoc test. **F**, Total nucleated cells (TNC) fold expansion of CD90+ HSC cultures for 7 days with 1μM Pep G, Pep D, or Pep S treatment; n = 10 independent cultures, *p = 0.05, one-way ANOVA with Dunnett’s post-hoc test. **G**, Percent CD90+ HSCs in culture after 7 days with 1μM Pep G, Pep D, or Pep S treatment; (n = 10 independent cultures; *p = 0.05, one-way ANOVA with Dunnett’s post-hoc test. **H**, Absolute CD90+ HSC counts in cultures after 7 days with 1μM Pep G, Pep D, or Pep S treatment; n = 10 independent cultures, *p = 0.05, one-way ANOVA with Dunnett’s post-hoc test.

To examine MA effects on HSCs, 10,000 phenotypic CD90^+^ HSCs were cultured for 7 days in serum-free StemSpan (supplemented with SCF, TPO, and FLT3L) under 5% O_2_ and treated with either 5nM small-molecule MAs or 1μM peptide MAs. Resorted CD90^+^ HSCs stained with MTR showed significantly increased mitochondrial length after culture with Cmp B, and to a lesser extent Cmp A, consistent with on-target mitofusin agonist activity (**Figure 2B**; **Supplementary Figures S2B, S2C**). Similarly, Pep S increased mitochondrial length relative to the Pep G control, with a weaker effect observed for Pep D (**Figure 2B**; **Supplementary Figure S2D**). Mean mitochondrial length in resorted HSCs was more consistent, but not significantly different from that of freshly isolated HSCs (**Figure 2B**). MFN2 expression was maintained in both freshly isolated and day 7-day cultured CD90^+^ HSCs, supporting the potential for on-target mitofusin agonist activity (**Supplementary Figure S2E**). Neither nucleated cell expansion, CD90^+^ HSC frequency, nor CD90^+^ HSC yield was altered by small-molecule MAs (**Figure 2C-2E**) or peptide MAs (**Figure 2F-H**) compared to DMSO controls, indicating that MAs do not expand HSC numbers *in vitro*.

HSC cultures were functionally assessed by xenotransplantation using 25% of each 7-day culture into sublethal irradiated NSG recipients, enabling standardized comparison of HSC activity across conditions. At 30 weeks post-transplantation, recipients of Cmp B-treated cultures showed ∼4.5-fold higher mean human chimerism than DMSO controls, whereas Cmp A did not significantly enhance engraftment (**Figure 3A, left**; **Supplementary Figures S3A, S3B**). Human PBMC lineage output was comparable between Cmp B and DMSO recipients, indicating that mitofusin agonist treatment did not induce lineage bias (**Figure 3A, right**). Consistently, Pep S-treated cultures produced ∼5.5-fold higher mean human chimerism than Pep G controls, whereas Pep D had no significant effect (**Figure 3B**; **Supplementary Figures S3C, S3D**).These data suggest that while unphosphorylated MAs (Cmp B and Peptide S) markedly enhance HSC potency during ex vivo culture, whereas PINK-1 phosphorylation-dependent MAs (Cmp A and Pep D) had no significant effect. Further analysis of primary recipients showed that bone marrow recipients receiving Cmp B- and Pep S-treated cultures contained prominent human CD90^+^ HSC compartments at 30 weeks post-transplantation while control cultures showed HSC compartment exhaustion, correlating with peripheral blood output and suggesting MA treatment supports long-term HSC engraftment (**Figures 3C, 3D**).To further test long-term repopulation, we performed secondary transplantation using primary recipient BM. At 30 weeks post-transplantation, both Cmp B and Peptide S culture secondary recipients consistently maintained ≥1% human chimerism (positive cutoff), while DMSO and Pep G culture recipients failed to repopulate upon secondary transplantation (**Figures 3E, 3G**). Furthermore, multilineage engraftment was observed in both Cmp B and Pep S secondary recipients (**Figures 3F, 3H**).

**Figure 3:**
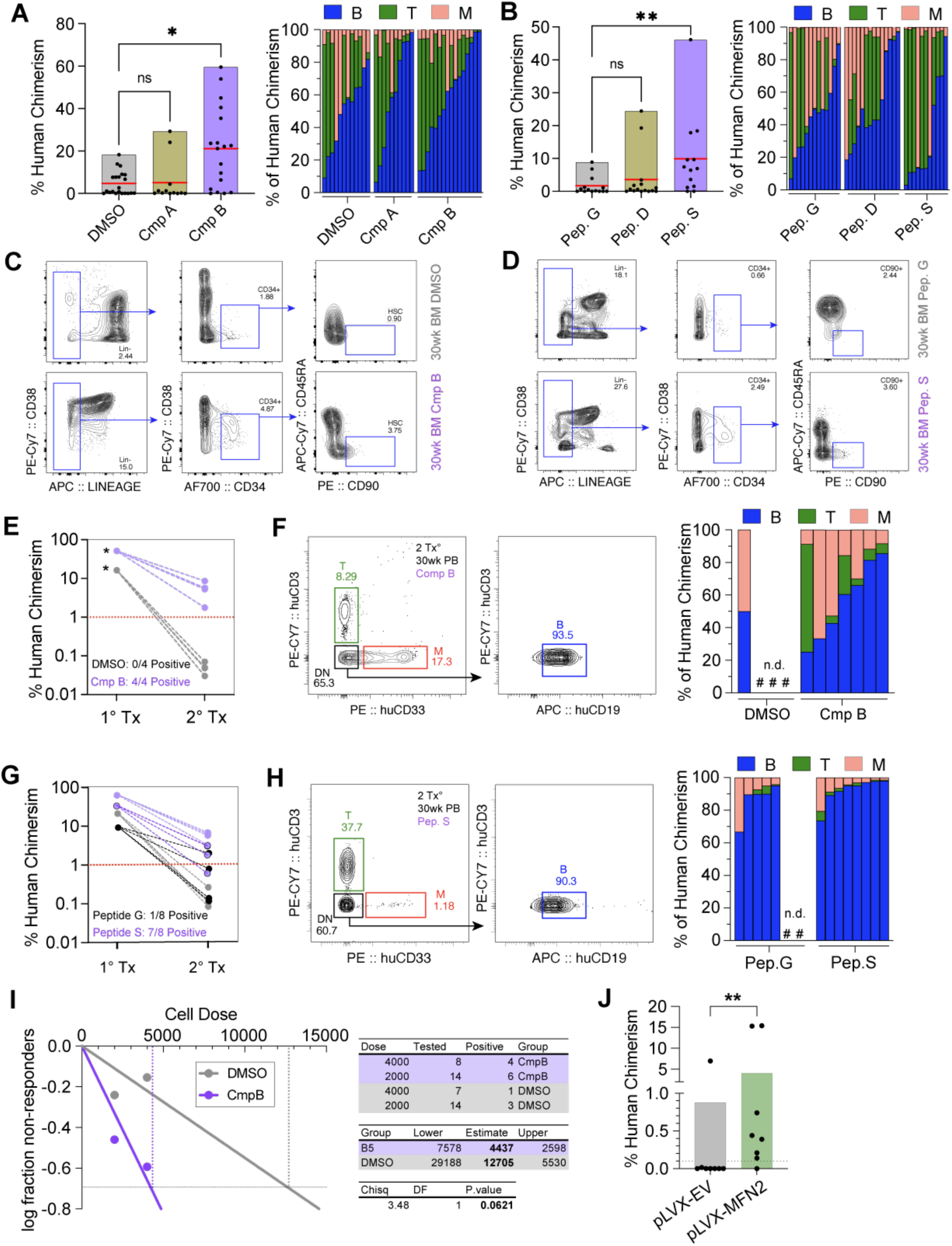
MAs improve long-term potency of CBU HSC *ex vivo* cultures. **A,** Primary human chimerism 30 weeks post-transplantation from recipients receiving 25% of CD90^+^ HSC cultures treated with DMSO, 5nM Cmp A, or 5nM Cmp B for 7 days; median (red bar), n≥15 recipients from five independent transplants; *p<0.05; Kruskal-Wallis test with Dunn’s post-hoc multiple comparisons test. **B,** Primary human chimerism 30 weeks post-transplantation from recipients receiving 25% of CD90^+^ HSC cultures treated with DMSO, 1μM Pep G, 1μM Pep D, or 1μM Pep S for 7 days; median (red bar), n≥15 recipients from five independent transplants; *p<0.05; Kruskal-Wallis test with Dunn’s post-hoc multiple comparisons test. **C,** Representative FACS phenotype of bone marrow 30 weeks post-transplantation from primary transplant recipients receiving cultures treated with DMSO or 5nM Cmp B. **D,** Representative FACS phenotype of bone marrow 30 weeks post-transplantation from primary transplant recipients receiving cultures treated with Pep G or Pep S. **E,** Secondary human chimerism 30 weeks post-transplantation from recipients receiving 1 × 10^6^ BM cells from DMSO or Cmp B primary NSG recipients. Line graphs show Δlog donor/competitor ratio from primary (1°) and secondary (2°) recipients. Threshold for positive chimerism is 1%; n = 4 one sample t-test; *p < 0.05. **F,** Representative PB FACS phenotype from secondary NSG recipients 30 weeks post-transplantation receiving DMSO or Cmp B cultures (left), and human PB lineage output showing myeloid (M, CD33^+^), B cell (B, CD19^+^) and T cell (T, CD3^+^) contributions from individual recipients (right). **G,** Secondary human chimerism 30 weeks post-transplantation from recipients receiving 1 × 10^6^ BM cells from Pep G or Pep S primary NSG recipients. Line graphs show Δlog donor/competitor ratio from primary (1°) and secondary (2°) recipients. Threshold for positive chimerism is 1%; n=8 recipients from 2 independent experiments, *p < 0.05, one sample t-test. **H,** Representative PB FACS phenotype from secondary NSG recipients 30 weeks post-transplantation receiving Pep G or Pep S cultures (left), and human PB lineage output showing myeloid (M, CD33^+^), B cell (B, CD19^+^) and T cell (T, CD3^+^) contributions from individual recipients (right). **I,** Limiting dilution analysis 15 weeks post-transplantation from recipients receiving high (4,000) and low (2,000) dose cell numbers of CD90^+^ HSCs cultured with DMSO or 5nM Cmp B for 7 days. The natural log fraction of non-responders (left) and estimated stem cell frequencies and statistical analysis table (right) is shown; n=8 high dose and n=14 for low dose, N=2 independent experiments, ELDA Poisson distribution; P<0.05; Pearson’s chi-square test. **J**, Primary human chimerism 8-weeks post-transplantation from recipients receiving 25% of CD90^+^ HSC cultures transduced with empty vector (pLVX-EV) or MFN2 lentivirus (pLVX-MFN2) for 72 hours; n = 8 recipients from two independent experiments; *p<0.05; Kruskal-Wallis test with Dunn’s post-hoc multiple comparisons test.

To more accurately quantify HSC frequency, we performed limiting dilution (LDA) transplantation assays. Cmp B-treated cultures increased the frequency of HSCs capable of generating >0.1% human chimerism at 15 weeks post-transplantation by ∼3-fold relative to DMSO culture controls, supporting preservation of long-term repopulating activity during *ex vivo* culture (**Figure 3I; Supplementary Figure 3E**). To confirm an MFN2-specific role, HSC where transduces lentivirus to overexpress MFN2. Transduction efficiency was 100%, as determined by IRES-GFP expression at 72 hours while preserving HSC phenotype (**Supplementary Figure S3F**). Expression of MFN2 mRNA was increased ∼3-fold relative to empty vector controls (**Supplementary Figures S3G**). At 8 weeks post-transplantation, MFN2-LV HSCs generated ∼4-fold higher human chimerism than empty vector-LV controls (**Figure 3J**). Taken together, these data strongly demonstrate elevated MFN2 expression and MA treatment improves long-term engraftment and multilineage potential UCB HSCs during ex vivo culture.

### Mitofusin agonists activate catabolic programs during *ex vivo* culture

MFN2 activity has been linked to changes in metabolism and bioenergetics.^23^ To explore the mechanistic basis of MAs, we first assessed its involvement in mitochondrial metabolic activity. Seahorse extracellular flux analysis showed no significant difference in mitochondrial respiration between CD34^+^ HSPC cultures treated 3 days with Cmp B or DMSO (**Supplementary Figure S4A, S4B**). Mitochondrial membrane potential, assessed by TMRM staining, and glucose uptake, measured by 2-NBDG incorporation, were comparable between CD90^+^ HSCs in Cmp B and DMSO cultures after 3 days, suggesting that MAs act independently of substrate metabolism. (**Supplementary Figure S4C-S4F**). MFN2-mediated mitochondrial fusion may also promote PARKIN-dependent mitophagy.^24^ To test whether mitofusin agonists alter this effect biochemically, MitoKeima-transfected HeLa cells were treated with MAs and assessed for mitochondrial acidification.^25^ Cmp B culture for 3 days did not alter mitochondrial acidification ratios relative to 3 hour CCCP control, indicating no detectable effect on mitophagic flux in HeLa cell lines. (**Supplementary Figure S4G, S4H**).

Next, we performed RNA-seq of resorted CD90^+^ HSCs after 7-day culture with Cmp B or DMSO. PCA analysis showed distinct gene expression patterns between treatments while differential expressed genes (DEG) analysis revealed 545 upregulated and 123 downregulated genes in Cmp B treated HSCs (**Figure 4A, 4B, Supplementary Table 1**). Notable Gene Ontology (GO) terms associated with upregulated genes included cytoplasmic translation, hematopoietic stem cell differentiation and cellular response to hypoxia while notable KEGG pathways included ribosomes and autophagy (**Figure 4C, Supplementary Table 2**). GSEA normalized enrichment scores were significant for pathways including cytoplasmic stress granule formation, cytosolic ribosome, and lysosomal vesicle biogenesis (**Figure 4D**). Interestingly, although not significant, numerous genes from the Eppert HSC GSEA gene set were upregulated in Cmp B treated CD90^+^ HSCs suggesting enhanced HSC identity (**Supplementary Figure S4I**).^26^ RNA-seq showed that *MFN2* expression was ∼200-fold higher than *MFN1* and was unchanged by Cmp B treatment, suggesting that MFN2 is the predominant mitofusin agonist target in CD90^+^ HSCs (**Supplementary Figure S4J**). Taken together, these data suggest an elevated stemness signature in CD90^+^ HSCs after *ex vivo* culture with MAs.^26^

**Figure 4:**
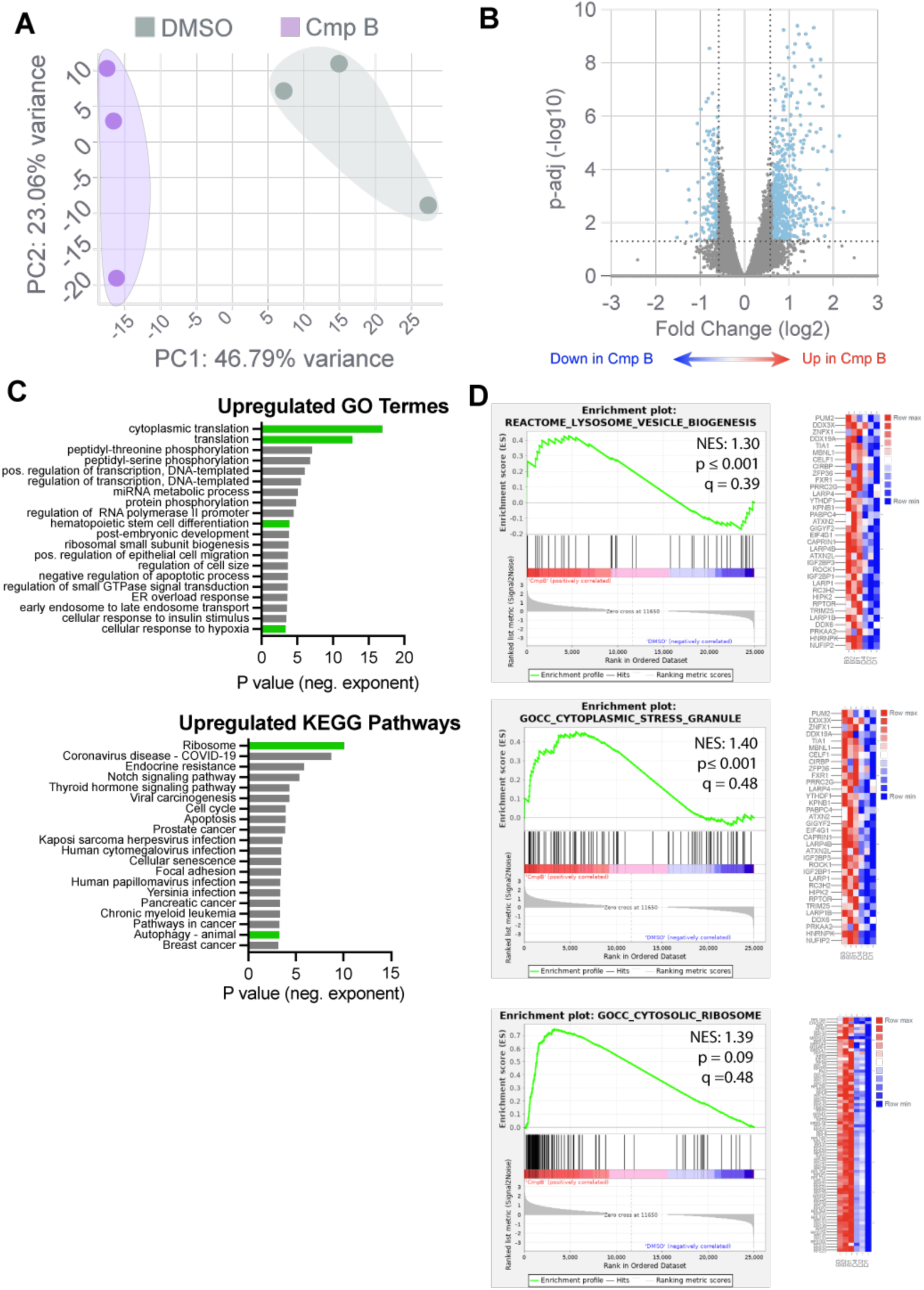
Upregulation of metabolic programs define HSCs cultured with MAs. **A**, PCA plot of RNA-seq data from resorted CD90^+^ HSCs after 7 days of culture with DMSO or 5nM Cmp B. **B**, Volcano plot depicting DGE from resorted CD90^+^ HSCs after 7 days of culture with DMSO or 5nM Cmp B; n=3 biological replicates, x-axis threshold ±50% fold change in expression, y-axis threshold p adjusted <0.05. **C**, Gene Ontology and KEGG pathway terms upregulated in CD90^+^ HSCs after 7 days of culture with 5nM Cmp B. Translation and autophagy terms are highlighted in green. **D**, GSEA plots of genes sets related to translation and autophagy upregulated in CD90^+^ HSCs after 7 days of culture with 5nM Cmp B. Z-score expression of genes in GSEA set are shown.

Ribosomal and lysosomal pathways, which regulate HSC function, were next evaluated in CD90^+^ HSCs following mitofusin agonist treatment. Analysis of global protein synthesis using O-propargyl-puromycin (OPP) staining of cultures showed a ∼30% reduction in translational activity with Cmp B treatment compared to control in CD90^+^ HSCs, although not to the level observed using the dual specificity phosphatase 2 (DUSP2) inhibitor, salubrinal, which specifically inhibits eIF2α–mediated translation (**Figure 5A, 5B**). Levels of phos-S51-eIF2α were ∼70% higher in CD90^+^ HSCs after Cmp B culture compared to control, indicating MA treatment mediates signaling to reduce protein synthesis, which is a critical feature of HSC maintenance (**Figures 5C-5D**).^27^ Protein aggregation, assessed by ProteoStat staining, was unchanged following Cmp B treatment compared with controls and in contrast to heat shock–treated cells, suggesting that MA treatment does not alter protein aggregation or chaperone-mediated autophagy pathways that have been implicated in HSC function. (**Figure 5D, Supplementary Figure S5A)**.^28,29^

**Figure 5:**
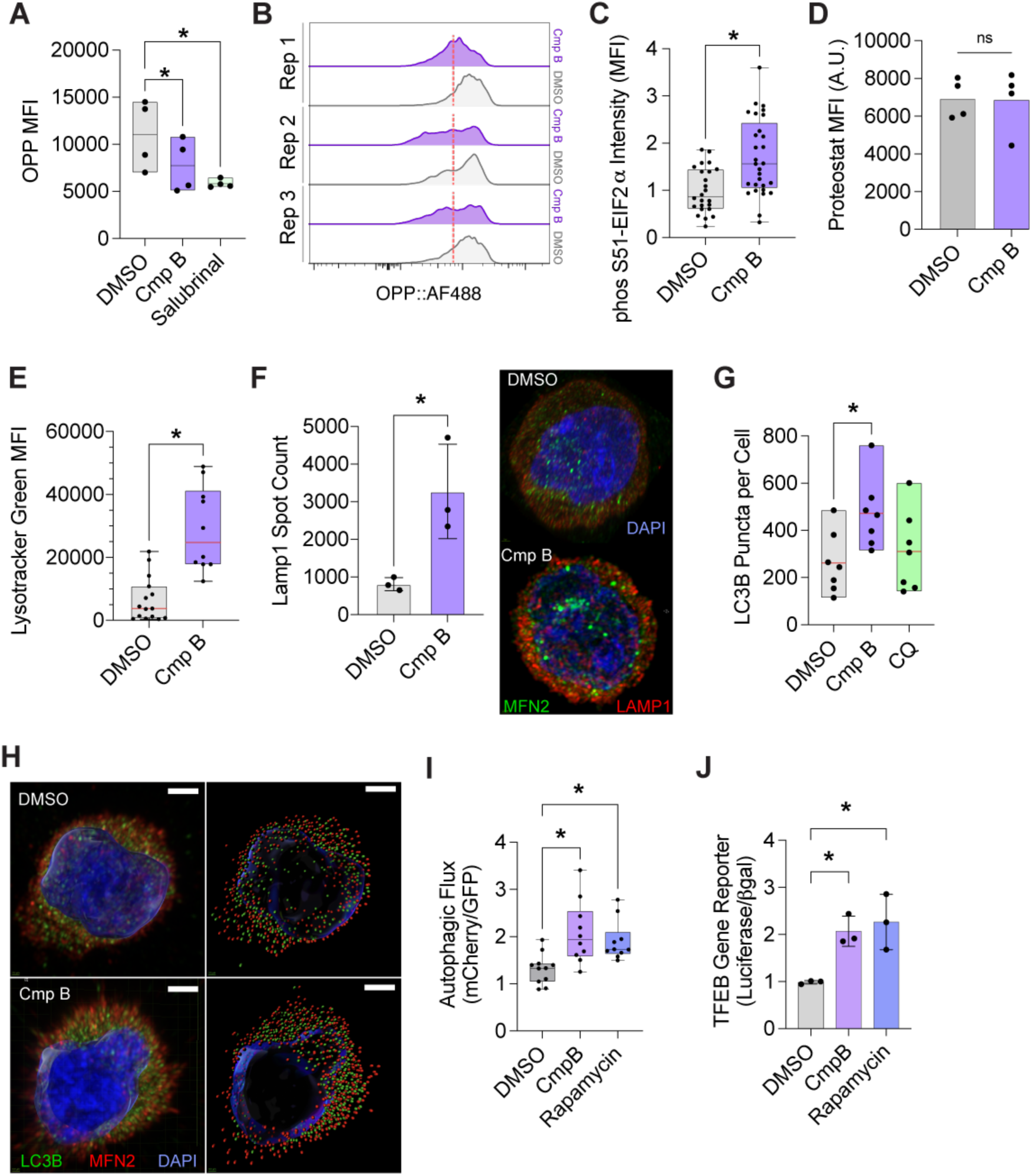
MA treatment reduces ribosome translation and stimulates autophagic flux in cultured CD90^+^ HSCs. **A**, Flow cytometric MFI of OP-Puro signal in CD90^+^ HSCs cultured for 3 days with DMSO, 5nM Cmp B, or 10μM salubrinal control (n = 4 biological replicates, *p < 0.05, one-way ANOVA). **B**, Representative histograms illustrating OPP signal in CD90^+^ HSCs expanded for 3d with DMSO or 5nM Cmp B. **C**, Image quantification of phospho-eIF2α intensity in resorted CD90^+^ HSCs cultured for 3 days with DMSO or 5nM Cmp B; n ≥ 20 cells from three biological experiments, *p < 0.05, student’s t-test). **D**, Flow cytometric MFI of ProteoStat signal in CD90^+^ HSCs cultured 3 days with DMSO or 5nM Cmp B; n = 4 biological replicates, *p < 0.05, two-tailed student t-test. **E**, Flow cytometric MFI of Lysotracker Green signal in CD90^+^ HSCs cultured for 3 days with DMSO or Cmp B; n= 16, *p < 0.05, student’s t-test. **F**, Spot count of LAMP1 IF staining (*left*) and representative super-resolution confocal images of MFN2 (green) and LAMP1 (red) staining (*right*) in resorted CD90^+^ HSCs cultured 3 days with DMSO or 5nM Cmp B; mean of n = 3 biological replicates, *p < 0.05, student’s t-test). **G**, Spot count of LC3B IF staining in resorted CD90^+^ HSCs cultured 3 days with DMSO, 5nM Cmp B or 50μM chloroquine (CQ) control; n= 7 cells from 2 biological replicates, *p < 0.05, one-way ANOVA. **H,** Super-resolution confocal micrographs (left) and Imaris spot analysis (right) of LC3B (green) and MFN2 (red) IF staining and DAPI nuclear stain (blue) in HSCs treated for 3d with DMSO or 5nM Cmp B. Scale bar is 1mm. **I,** Image quantification of autophagic flux reporter ratio (mCherry/LC3-GFP) in CD34^+^ cells transfected with FUW mCherry-GFP-LC3 plasmid followed by culture for 3 days in DMSO, 5nM Cmp B, or 10μM rapamycin; n = 10 cells from two biological replicates, *p < 0.05, two-tailed student t-test. **J**, Luciferase gene reporter activity (luciferase/βGal) in CD34^+^ cells transfected with pLminP-RE28 and pSV-βGal plasmids followed by culture for 3 days in DMSO, 5nM Cmp B, or 10μM rapamycin; n = 3 biological replicates, *p < 0.05, two-tailed student t-test.

Next, we assessed lysosome content and autophagic flux in MA-treated cultures. Lysotracker staining showed >2-fold increase in lysosomal content in CD90^+^ cells cultured in Cmp B compared to control (**Figures 5E**). To further assess autophagic content, LAMP1 and LC3B puncta were quantified by SCRM. Cmp B-treated CD90^+^ HSCs showed a ≥3-fold increase in LAMP1^+^ lysosomal puncta and a ≥2-fold increase in LC3B^+^ autophagosome puncta compared with controls. (**Figures 5F-5H**). While Cmp B treatment did not significantly alter the total number of MFN2 puncta compared with DMSO control, MFN2–LAMP1 colocalization was significantly reduced following Cmp B treatment suggesting decreased spatial association between MFN2^+^ structures and lysosomes (**Supplementary Figure S5B, S5C**). To confirm the effect of MAs on autophagic flux, the mCherry-GFP-LC3B reporter was used by exploiting the differential pH sensitivity of GFP and mCherry.^30,31^ Upon autophagosome fusion with lysosomes, GFP fluorescence is quenched in the acidic autolysosome, whereas mCherry fluorescence is retained, increasing the mCherry/GFP ratio. Thus, a higher mCherry/GFP ratio reflects increased autophagic flux. SCRM analysis of CD34^+^ HSPCs transfected with mCherry-GFP-LC3B reporter and cultured with Cmp B for 3 days showed increased mCherry/GFP ratio by ∼2-fold relative to DMSO and comparable to rapamycin, indicating enhanced autophagic activity in HSPCs following MA treatment (**Figure 5I**).

Transcription Factor EB (TFEB) is a master transcriptional regulator of lysosomal biogenesis that promotes expression of genes involved in lysosome formation, acidification, and degradative function.^32,33^ TFEB nuclear translocation upregulates a coordinated lysosomal gene expression network, thereby increasing lysosomal capacity and autophagic clearance and consistent with increased lysosomal biogenesis and activation of autophagy–lysosome pathways. Therefore, TFEB-dependent lysosomal biogenesis was evaluated using complementary gene reporter and transcriptomic approaches.^34^ In CD34^+^ cells, Cmp B treatment significantly increased TFEB reporter activity in 3-day cultures relative to DMSO controls, reaching levels comparable to rapamycin and consistent with activation of TFEB-dependent transcription (**Figure 5J**). To determine whether this reporter activity was reflected in the transcriptional response of HSCs, DEGs from Cmp B-treated HSCs (**Figure 4**) were cross-referenced with curated TFEB target genes in the Gene Transcription Regulation Database.^35^ A significant enrichment in putative TFEB-regulated genes were identified, including 11% of upregulated and 4% of downregulated DEGs, supporting TFEB activation as a mechanism by which MA treatment enhances lysosomal biogenesis and autophagic flux. (**Supplementary Figure 5D, Supplementary Table 3**). Taken together, these data indicate MA treatment induces a catabolic program that attenuates global protein synthesis, activates autophagy, and increases lysosomal content to support long-term repopulation potential of HSCs during *ex vivo* culture.

### MFN2 interacts with MTOR to attenuate downstream anabolic signaling

Although MA treatment clearly affected protein synthesis and autophagy, how MAs modulates these pathways remained unclear. To explore regulatory networks of differentially expressed genes in Cmp B-treated HSCs, we employed the STRING Interactome in Network Analyst.^36^ Clustering of enriched GSEA gene sets for ribosomes, autophagy, and stress granules revealed a central node involving mammalian target of rapamycin (MTOR) signaling (**Supplementary Figure 5E**). Because MTOR modulates protein synthesis, autophagy, and stem cell activity, we hypothesized that MA treatment alters MTOR signaling in HSC cultures, thus explaining the mechanism of improved *in vivo* engraftment.

FACS analysis of short-term CD90^+^ HSC cultures showed MTOR expression was not altered with Cmp B treatment (**Figure 6A, Supplementary Figure S6A**). However, activated MTOR levels using phos-S2448-MTOR specific staining were reduced ∼50% in CD90^+^ HSCs after Cmp B culture (**Figure 6B, Supplementary Figure 6B**). Phosphorylated RPS6 (phospho-S6) protein levels, a downstream target of MTOR kinase activity required for enhanced translation and global protein synthesis,^37^ were reduced 22% in CD90^+^ HSCs cultured with Cmp B compared to DMSO and were further reduced in rapamycin treatment control cultures (**Figure 6C, Supplementary Figure S6C**). Since upstream MTOR signaling can occur rapidly, mitofusin agonist effects were assessed in HeLa cells after 1 hour of treatment.^38^ Cmp B and rapamycin did not alter total MTOR or phospho-AKT levels, indicating that upstream AKT–MTOR signaling was unchanged. However, Cmp B reduced phospho-S6 levels by ∼25% relative to control, although less robustly than rapamycin, suggesting MA treatment partially inhibits downstream MTOR activity (**Figures 6D-6F**).

**Figure 6:**
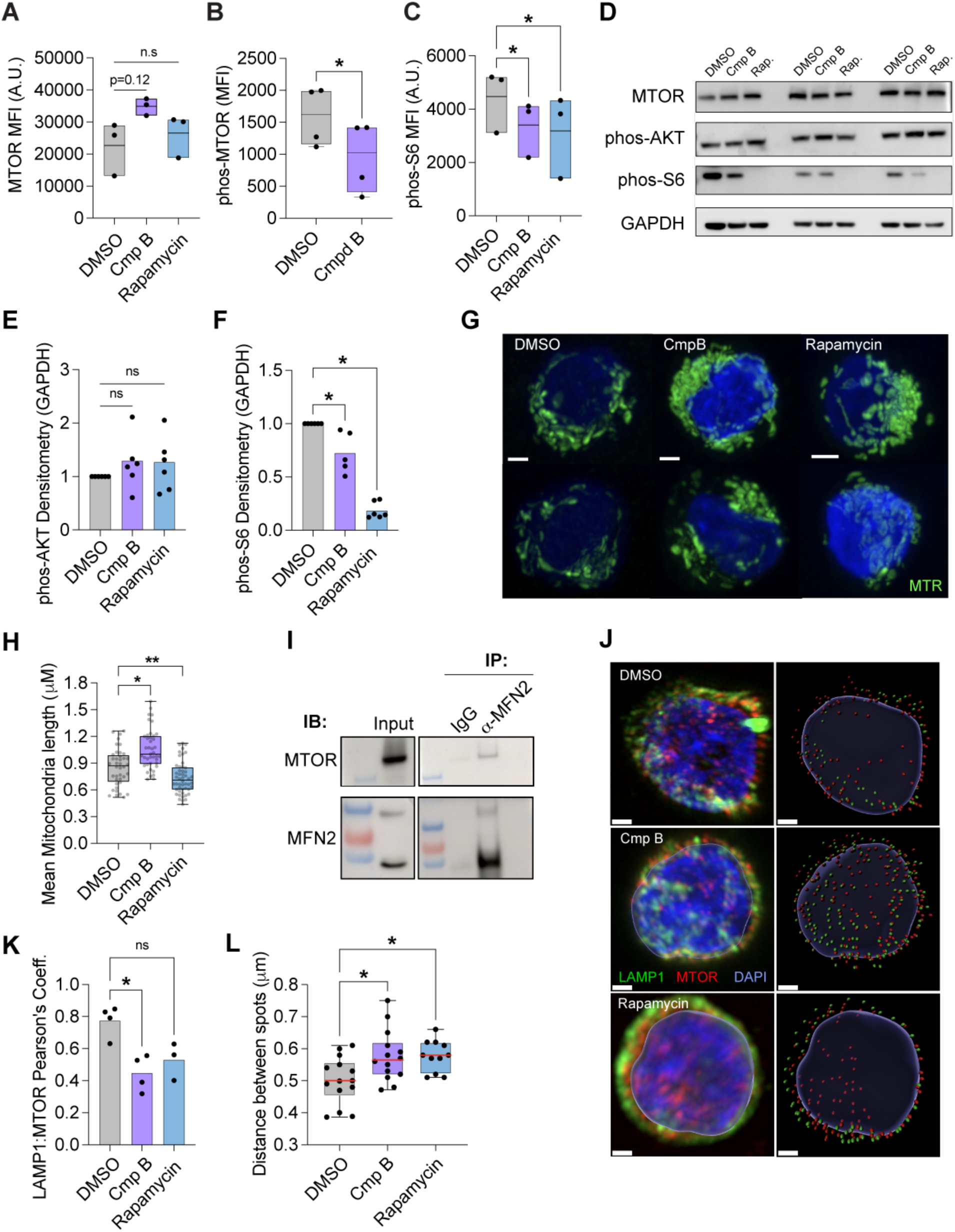
MFN2 interacts with MTOR and correlates with reduced downstream signaling in cultured CBU HSCs. **A**, Flow cytometric MFI of MTOR intensity in CD90^+^ HSCs cultured for 3 days with DMSO, 5nM Cmp B, or 10μM rapamycin; n=3 independent experiments, *p < 0.05, one-way ANOVA with Dunnett’s post-hoc test. **B**, Flow cytometric MFI of phos-S2448 mTOR intensity in CD90^+^ HSCs cultured for 3 days with DMSO or 5nM Cmp B; n=4 independent experiments, \**p* < 0.05, student’s *t*-test. **C**, Flow cytometric MFI of phos-S6 intensity in CD90^+^ HSCs cultured for 3 days with DMSO, 5nM Cmp B, or 10μM rapamycin; n= 3, \**p* < 0.05, one-way ANOVA with Dunnett’s post-hoc test. **D,** Representative western blots from HeLa cells treated for 1h with DMSO, 5nM Cmp B or 10uM rapamycin. Blots show lysates from three independent experiments. **E,** Densitometry of phospho-AKT in HeLa cell lysates cultured for 1h with DMSO, 5nM Cmp B or 10uM rapamycin. Protein levels were normalized with GAPDH loading control protein; n=6 independent experiments, \**p >* 0.05, one-way ANOVA with Dunnett’s post-hoc test. **F**, Densitometry of phospho-S6 in HeLa cell lysates cultured for 1h with DMSO, 5nM Cmp B or 10uM rapamycin. Protein levels were normalized with GAPDH loading control protein; n=6 independent experiments, \**p >* 0.05, one-way ANOVA with Dunnett’s post-hoc test. **G**, Representative immunofluorescent images of fresh CBU CD90^+^ HSCs, or CD90^+^ HSCs cultured for 3 days with DMSO, 5nM Cmp B or 10μM rapamycin followed by staining with MTR (green) and DAPI (blue). **H,** Quantification of mitochondrial lengths from freshly sorted CD90^+^ HSCs, or CD90+ HSCs cultured for 3 days with either DMSO, 5nM Cmp B, or 200nM rapamycin stained and with MTR; average of ≥100 mitochondria from n ≥ 12 cells from two independent experiments, \**p* < 0.05, one-way ANOVA with Dunnett’s post-hoc test. **I**, Representative western blot of endogenous MFN2 immunoprecipitation with MTOR protein in 293T cell lysates. **J**, Super-resolution confocal micrographs (left) and Imaris spot analysis (right) of LAMP1 (green), mTOR (red) in resorted CD90^+^ HSCs cultured for 3d with DMSO, 5nM Cmp B or 10μM rapamycin. Scale bar is 1mm. **K**, Pearson’s correlation coefficients of MTOR and LAMP1 confocal microscopy in resorted CD90^+^ HSCs cultured for 3 days with DMSO, 5nM Cmp B or 10uM rapamycin. Data derived from the average of n= 4 fields of cells from 3 independent experiments; \**p* < 0.05, one-way ANOVA with Dunnett’s post-hoc test. **L**, Imaris spot distance between LAMP1 and MTOR spots in resorted CD90^+^ HSCs cultured for 3 days in DMSO, 5nM Cmp B, or 10μM rapamycin; data derived n = 10-12 individual cells across 3 independent experiments; distance calculated using the 3-nearest neighbor method, \**p* = 0.05, one-way ANOVA with Dunnett’s post-hoc test.

Changes in MTOR signaling affect mitochondrial function and morphology, including alterations in mitochondrial length.^39^ CD90^+^ HSCs treated with Cmp B or rapamycin for 3 days and stained with MTR showed Cmp B increased mitochondrial length by ∼44% compared with DMSO, whereas rapamycin had no effect, indicating that mitochondrial elongation was specific to MA treatment (**Figures 6G, 6H**). Since MFN2 has previously been shown to interact with the MTORC2 complex and given our findings, we surmised MFN2 may interact with MTOR to regulate its activity.^40^ To test this, we performed immunoprecipitation in cell lines. MFN2 pulldown in 293T cell lysates showed clear precipitation of MTOR protein, confirming potential interaction (**Figure 6I**). Further, overexpression of 6x-His tagged MFN2 followed by pulldown further verified interaction between overexpressed MFN2 and endogenous MTOR protein, but not in empty vector controls (**Supplementary Figure 6E, 6F**). Protein docking simulations using crystal structures for human the mTORC1 complex and MFN2 revealed a significant interaction between the FKBP12 rapamycin binding (FRB) domain of MTOR and the HR1 and HR2 domains of MFN2 (**Supplementary Figure 6D**) further substantiating interaction potential between MTOR and MFN2.^41,42^ Finally, we surmised that if MTOR physically associates with mitochondrial MFN2, then colocalization experiments would show physical disassociation of MTOR from lysosomes, which is the site of MTOR activation.^43^ To test this, SRCM between LAMP1 and MFN2 was used to estimate distances. Confocal micrographs showed a significant reduction in MTOR and LAMP1 colocalization in CD90^+^ cells cultured with either Cmp B or rapamycin (**Figure 6J, 6K**). Furthermore, physical distances between MTOR and LAMP1 was increased with either Cmp B or rapamycin treatment (**Figure 6L**). In xenotransplantation assays, both CD90^+^ HSCs and CD34^+^ HSPCs cultured with Cmp B showed enhanced human engraftment compared with controls (**Supplementary Figures S6G, S6H**). In contrast, CD90+ HSC rapamycin-treated cultures showed reduced engraftment relative to DMSO while CD34+ HSCP rapamycin treated cultures were increase ∼2-fold to control. These data suggest that mitofusin agonists and MTOR inhibition support functional repopulating activity of cultured HSCs and HSPCs. Taken together, these data suggest MA treatment facilitates MFN2 sequestration of MTOR to attenuate downstream protein synthesis and autophagy programs that maintain *in vivo* HSC repopulation potential during *ex vivo* culture.

### MFN2 is partially required to regulate MTOR-mediated proteostasis, autophagy and lysosomal programs

To confirm the role of MFN2 in Cmp B-mediated MTOR regulation, we generated a MFN2 knockout (KO) 293FT cell line (**Figure 7A, 7B**). In wild-type (WT 293) cells expressing Mito-dsRed, Cmp B treatment increased mitochondrial length by ∼44%, and rapamycin mirrored mitochondria elongation effects (**Figure 7C, 7D**), corroborating studies showing MTOR inhibition enhances fusion in some cell types.^39,44^. By contrast, MFN2 KO cells displayed punctate, toroidal-shaped mitochondria (∼30% shorter than WT).^45,46^ Notably, both Cmp B and rapamycin partially restored MFN2 KO mitochondrial length to WT levels, likely via MFN1-mediated fusion activity, consistent with reports that MAs rescue mitochondrial dysmorphology in MFN2 KO MEFs due to MFN1 expression that cannot be rescued in MFN1/MFN2 double KO MEFs.

**Figure 7:**
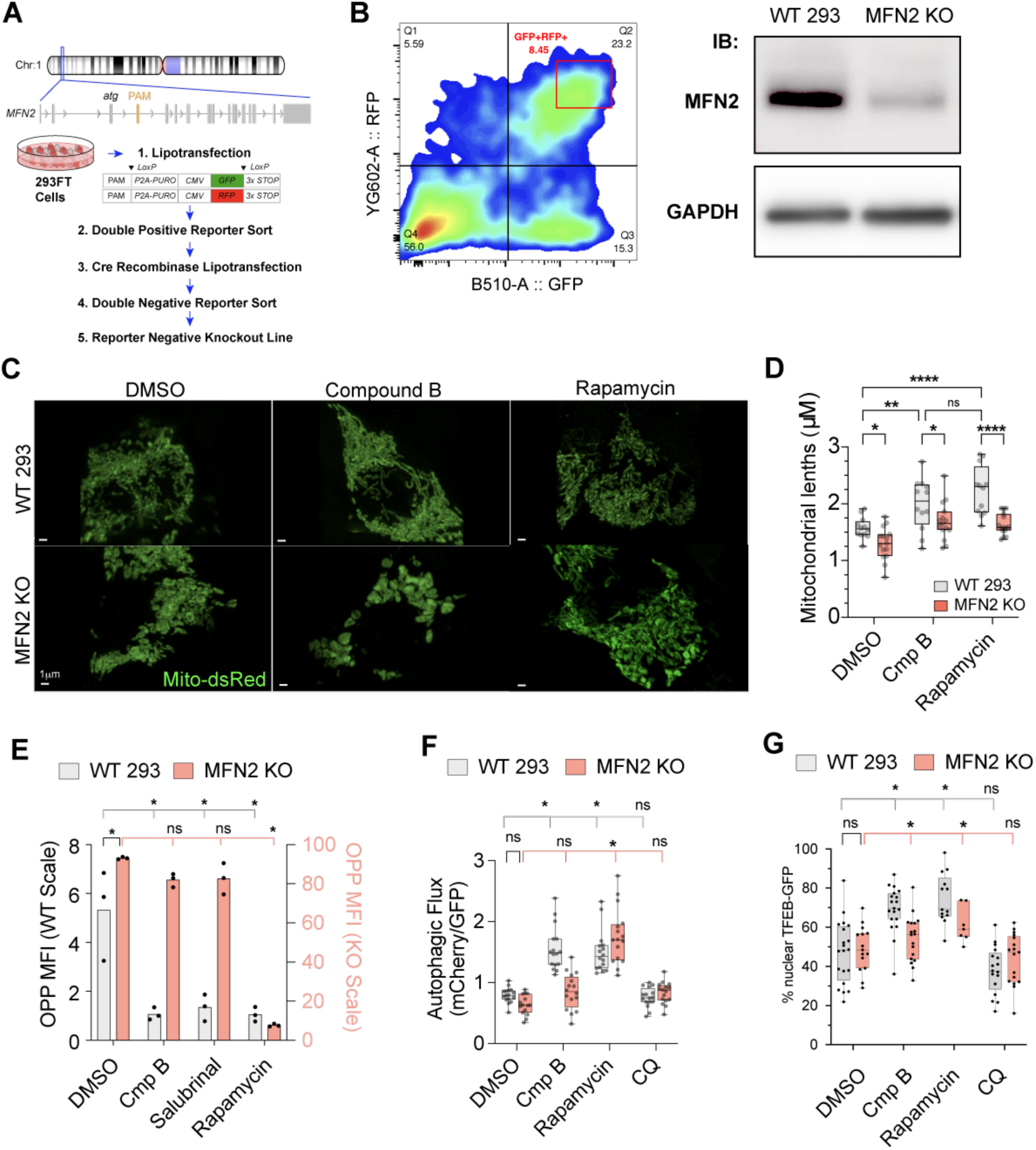
Loss of MFN2 inhibits MA attenuation of global protein translation and lysosomal biogenesis. **A**, Schematic representation of genetic editing strategy employed to produce MFN2 KO 293FT cell line. **B**, Representative gating strategy for double positive cells transfected sgRNAs with GFP or RFP reporter indicating disruption of MFN2 expression in both alleles (*left*). Representative western blot for MFN2 and GAPDH protein levels in WT 293 and MFN2 KO 293 cell lysates (*right*). **C**, Representative images of mitochondria morphology in wild type (WT 293) and MFN2 KO 293 cells transfected with Mito-dsRed reporter and cultured with DMSO, 5nM Cmp B, or 200nM rapamycin for 48h. **D**, Quantification of mitochondria lengths in resorted WT and MFN2 KO cells cultured 3 days with DMSO, 5n Cmp B or 200nM rapamycin. (≥ 10 cells and ≥ 100 mitochondria across 3 independent experiments); *p<0.05, two-Way ANOVA with Tukey’s multiple comparison test. **E**, Quantification of OP-Puro levels in WT 293 and MFN2 KO 293 cells cultured for 3 days with DMSO, 5nM Cmp B, 10μM salubrinal, or 200nM rapamycin; n= 3, \**p* < 0.05, one-way ANOVA with Dunnett’s post-hoc test. **F,** Image quantification of autophagic flux reporter ratio (mCherry/LC3-GFP) in 293 WT and MFN2 KO 293 cells transfected with FUW mCherry-GFP-LC3 plasmid and then cultured for 3 days with DMSO, 5nM Cmp B, 10μM rapamycin, or 50μM chloroquine (CQ); n=16 fields of cells from 3 biological replicates, *p < 0.05, two-Way ANOVA with Tukey’s multiple comparison test. **G**, Nuclear TFEB-GFP signaling in WT 293 and MFN2 KO 293 cells transfected with TFEB-GFP followed by 3-day treatment with DMSO, 5nM Cmp B, 10μM rapamycin, 50μM chloroquine (CQ), or in combination; n≥ 25 cells from 3 independent experiments, *p<0.05, two-Way ANOVA with Tukey’s multiple comparison test.

OPP incorporation was increased ∼1.5-fold in MFN2 KO 293 cells relative to WT 293 controls, consistent with elevated protein synthesis following loss of MFN2 (**Figure 7E**). In WT 293 cells, Cmp B, salubrinal, and rapamycin reduced OPP signal relative to DMSO, whereas MFN2 KO 293 cells maintained high OPP levels after B5 and salubrinal treatment. Rapamycin markedly reduced OPP incorporation in MFN2 KO cells, suggesting that increased protein synthesis caused by MFN2 loss remains sensitive to MTOR inhibition. These results indicate that loss of MFN2 leads to dysregulated protein synthesis.

To evaluate whether MFN2 contributes for MTOR signaling, we measured autophagic flux and TFEB localization. Autophagic flux as measured by transfection with mCherry-GFP-LC3B reporter plasmid was comparable between WT 293 and MFN2 KO 293 cells under DMSO conditions, indicating that loss of MFN2 alone did not significantly alter basal flux (**Figure 7F**). Cmp B treatment increased autophagic flux in WT 293 cells, but this response was attenuated in MFN2 KO cells, supporting an MFN2-dependent effect of mitofusin agonist treatment. In contrast, rapamycin increased autophagic flux in both WT and MFN2 KO cells, whereas chloroquine had no significant effect on the flux ratio relative to baseline levels. To further substantiate a role for MAs in regulating lysosomal biogenesis, TFEB-GFP overexpression was performed in WT 293 and MFN2 KO 293 cells to visualized TFEB nuclear translocation.^47^ Nuclear TFEB-GFP was significantly increased by Cmp B and rapamycin treatment in WT cells, correlating with activation of TFEB-dependent lysosomal biogenesis. Cmp B treatment response was significantly attenuated in MFN2 KO cells, suggesting that MA-induced TFEB activation is at least partially MFN2-dependent. In contrast, rapamycin increased TFEB nuclear localization in both WT 293 and MFN2 KO cells, while chloroquine treatment did not suppress TFEB-GFP localization significantly compared to DMSO control. These data support differential regulation of TFEB activity and lysosomal pathways by MFN2 and MA treatment. Collectively, these findings indicate MFN2 partially regulates MTOR-dependent processes, including protein synthesis, autophagy, and lysosomal activity.

## Discussion

### Mitofusin agonists preserve long-term HSC repopulating activity during ex vivo culture

This study identifies mitofusin agonists (MAs) as a strategy to preserve the long-term engraftment potential of human HSCs during *ex vivo* culture. Chemically distinct MFN2 agonists, including the small molecule Cmp B and peptide mimetic Pep S, enhanced mitochondrial elongation and significantly increased long-term human chimerism after transplantation. The concordant activity of these structurally distinct agonists reduces the likelihood that improved HSC function reflects compound-specific off-target effects and supports MFN2 activation as the principal mechanism. Notably, MA treatment enhanced functional repopulating activity without expanding phenotypic HSC numbers, indicating that MAs primarily preserve HSC potency rather than increasing immunophenotypic HSC abundance.

### Fusion-competent MFN2 as a regulator of MTOR signaling

Mechanistically, our findings support a model in which MAs stabilize an open, fusion-competent conformation of MFN2 that promotes mitochondrial fusion while enabling MFN2-dependent attenuation of MTOR signaling. Immunoprecipitation, SRCM, and structural modeling suggest that MFN2 can associate with MTOR, potentially through interaction between the MFN2 HR2 domain and the MTOR FRB domain.^41,42^ Although the direct molecular interface requires further validation, these data support a model in which an MA-stabilized MFN2 protein complex sequesters MTOR away from RHEB-dependent lysosomal activation sites, thereby reducing downstream MTOR signaling. This interpretation is consistent with prior evidence that the MFN2 HR1 domain is capable of interaction with RICTOR to localize MTORC2 to mitochondrial membranes and suppress AKT signaling, supporting a broader role for MFN2 as a negative regulator of MTOR pathway activity.^40^

The lack of effect observed with PINK1 phosphorylation-dependent MAs, including Cmp A and Pep D, further suggests that the HSC-preserving activity of MAs depends on fusion-competent MFN2 rather than MFN2 functions associated with PINK1-regulated mitophagy. Phosphorylation of MFN2 at S378 by PINK1 has been linked to disruption of fusion and promotion of mitophagy.^48^ Since S378-dependent agonists did not significantly improve HSC repopulating activity in our study, our findings are consistent with prior studies linking quality control of mitochondrial via mitophagy to HSC maintenance.^49^ These data indicate that maintenance of long-term HSC function is preferentially associated with unphosphorylated, fusion-competent MFN2 and not fusion-inhibited phosphor-S378 mitochondria. Thus, unphosphorylated MAs preserve HSC potency by promoting fusion among healthy mitochondrion to maintain network integrity.

### Relationship to MTOR regulation of HSC fate

MTOR signaling is a central regulator of HSC differentiation, metabolism, and stress responses. Genetic studies using Mx1-Cre–mediated deletion of Raptor or Rictor have demonstrated that MTORC1 primarily regulates HSC differentiation, whereas MTORC2 has context-dependent roles in HSC development and function.^50–52^ Consistent with this biology, loss of the MTOR activator RHEB expands the HSC compartment but impairs downstream hematopoiesis, while MTORC1 activation promotes HSC differentiation that can be mitigated by rapamycin.^53–56^ Prior studies also showed that rapamycin treatment during short-term *ex vivo* culture of cord blood CD34^+^ cells can improve human chimerism after transplantation, supporting the concept that restrained MTOR activity preserves HSC potency.^54,57,58^ Our data extend these observations by identifying MFN2 as an endogenous regulator capable of attenuating MTOR signaling during HSC culture. Unlike rapamycin, which directly inhibits MTOR kinase activity, MAs appear to act through MFN2 conformation and localization. Cmp B promoted mitochondrial elongation, reduced protein translation, increased TFEB-associated lysosomal biogenesis, and enhanced autophagic flux. These findings support a model in which MA-stabilized MFN2 simultaneously promotes mitochondrial fusion and limits MTOR activation at lysosomal compartments, thereby preserving long-term HSC repopulating activity during culture.

### Autophagy, proteostasis, and translational restraint in HSC maintenance

Culture-associated stressors, including growth factor exposure, nutrient availability, and oxygen tension activate pathways that can increase protein synthesis, impair proteostasis, and reduce HSC potency.^59^ Autophagy is essential for HSC maintenance, and loss of autophagy genes has been shown to increase amino acid transporter expression, activate MTOR, and promote anabolic programs that compromise HSC function.^60–63^ Similarly, low global protein synthesis is required for normal HSC activity, while elevated translation and accumulation of misfolded proteins contribute to proteotoxic stress and premature HSC exhaustion.^28,64^ SIRT7-mediated mitochondrial unfolded protein response checkpoints further support the importance of proteostasis control in preventing HSCs in the context of aging.^65^

Consistent with these principles, MA-treated HSC cultures showed reduced global protein synthesis and downstream translation-associated signaling as well as increased autophagy–lysosome activity. Although ribosomal protein genes were increased after MA culture, the relationship between ribosomal gene expression and reduced protein synthesis may appear paradoxical but may reflect a poised HSC state wherein MAs may place HSCs in a “translation-ready but translationally restrained” state. This state may preserve stem cell potency during culture while maintaining the capacity for subsequent hematopoietic differentiation. Further studies will be needed to define the relative contributions of MTORC1 and MTORC2 to this response and to determine whether MFN2 regulates distinct MTOR complexes in HSCs.

### Translational implications for cord blood transplantation and gene therapy

Limited numbers of functional HSCs remain a major barrier to broader clinical use of cord blood units for transplantation. Current guidelines recommend minimum CD34^+^ and total nucleated cell thresholds for transplantation consideration, yet only a minority of banked cord blood units meet these criteria for a typical adult recipient.^66,67^ Maintaining long-term engraftment potential is also a major challenge for ex vivo gene transfer and genome editing approaches, where culture duration can erode HSC potency despite technical advances in genetic modification.^68^ The ability of MAs to preserve functional HSC activity during culture therefore has potential relevance for both cord blood transplantation and autologous or allogeneic gene therapy platforms.

In summary, these findings identify fusion-competent MFN2 as a previously unrecognized regulator of MTOR signaling and HSC potency during *ex vivo* culture. MAs preserve long-term repopulating activity by promoting MFN2-dependent mitochondrial fusion while restraining MTOR-associated protein synthesis and enhancing TFEB-linked lysosomal biogenesis and autophagic flux. Because the mechanisms of MAs appear distinct from existing HSC expansion strategies such as UM171 and nicotinamide derivatives, combinatorial approaches may provide a rational path to expand clinically relevant sources of HSC while preserving long-term engraftment potential. Future studies should define the quantitative effects of MAs on functional HSC frequency, determine optimal combinations with established expansion platforms, and assess whether this strategy can improve the clinical performance of cord blood and gene-modified HSC products.

## Acknowledgements

This work was supported by grants NIH R01 HL155574 (L.L.L.) and the AABB National Blood Foundation Early Career Grant. The content is solely the responsibility of the authors and does not necessarily represent the official views of the NIH. We would like to thank Jessica Freedman for assistance with experiments, Drs. Mohan Narla, Karina Yazdanbakhsh, Xuili An, and Patricia Shi for inciteful discussion.

## Authorship Contributions

G.C. and A.B. performed most experiments, contributed to the concept, and co-wrote the manuscript with L.L.L. D.J. and Z. Liang performed SCRM and transplantation experiments in Figures 2 & 3. J.F., Z. Leontiou, and E.L. assisted with CBU experiments in Figures 5 & 6. E.L. and K.L. provided support for multiple methods, PB and BM analyses and preparation of human CB cells. D.M. designed and validated MFN2 gene editing of MFN2 in Figure 7. N.N. and Y. C. helped co-write the manuscript and methods with L.L.L. C.D.H and H.-W.S. provided concept and guidance and co-wrote the manuscript. L.L.L. conceived of and designed the study and wrote the manuscript.

## Data Availability

Data and materials availability 26 Sequencing files and metadata. All data needed to evaluate the presented conclusions are in the main text and Supplemental Materials. All other data/materials will be distributed by authors upon request.

