## Supplemental Figures for "Mitofusin agonists improve the long-term repopulating activity of human cord blood HSCs after *ex vivo* culture"

**Supplementary Figure 1**

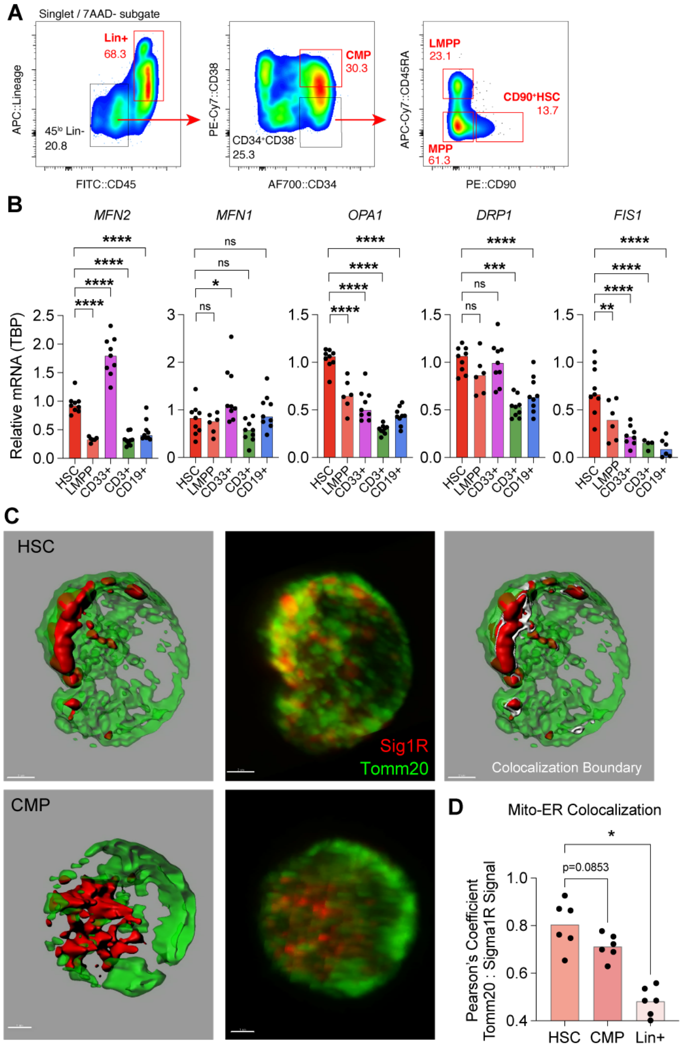

**Supplementary Figure 1: CBU population gating and mitochondria and ER imaging.**

**A**, Representative FACS gating strategy used to identify phenotypic CBU cell populations.

**B,** Relative mRNA expression of mitochondrial dynamic genes in HSC, LMPP, CD33+(myeloid), CD3+ (T cell), and CD19+ (B cell) cells. mRNA levels were normalized with TBP housekeeping gene; n=9 from 3 independent experiments, *$p < 0.05$, one-way ANOVA with Dunnett's post-hoc test.

**C**, Representative IF images of the mitochondrial marker TOMM20 (green) and the ER marker S1R (red) in CBU HSCs and CMPs. Imaris surface rendering of each signal is shown. White surfaces indicate colocalized surface boundaries. Scale bar is 1μm.

**D**, Pearson's colocalization analysis of S1R voxels colocalized with TOMM20 voxels in CBU HSCs, CMPs and Lin+ cells; n = 6 cells from 2 independent experiments, *$p < 0.05$, one-way ANOVA with Dunnett's post-hoc test.

### Supplementary Figure 2

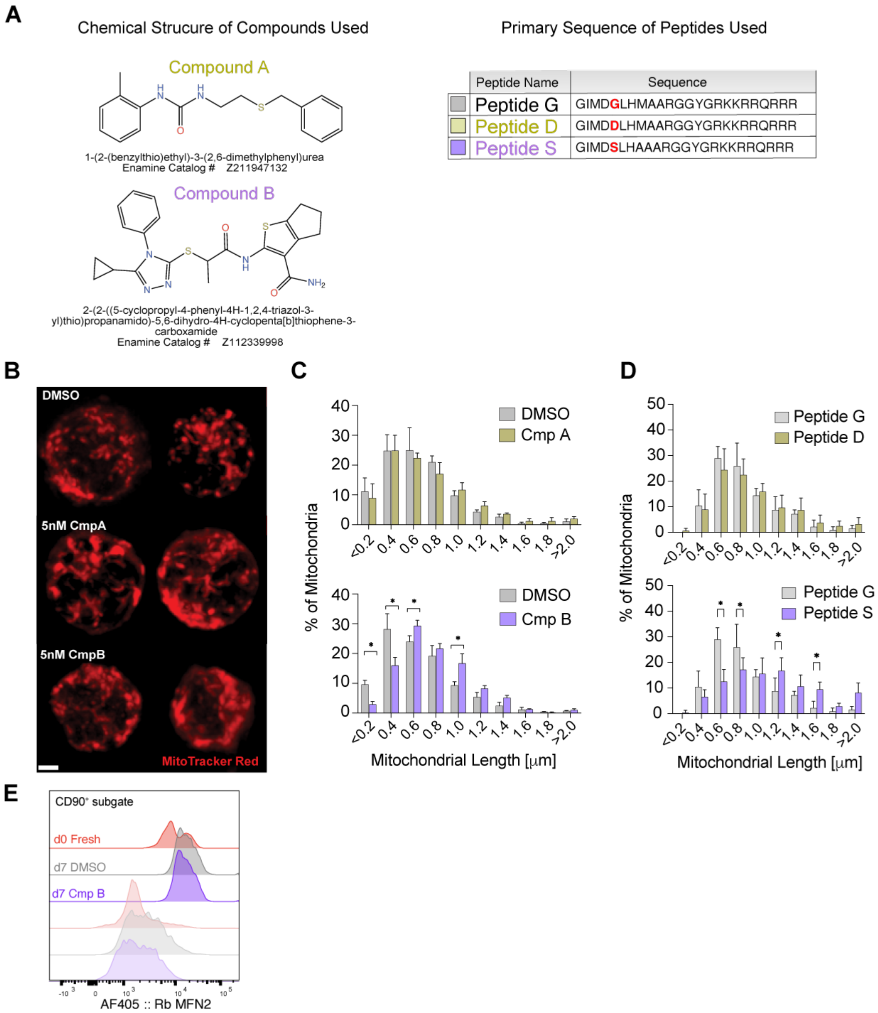

| | Sample Name | Mean : 405-F-A |
|---|---|---|
| | CD90+ Day 0 (Fresh) - anti MFN2 | 11322 |
| | CD90+ Day 7 DMSO - anti MFN2 | 19499 |
| | CD90+ Day 7 5nM CmpB - anti MF | 17955 |
| | CD90+ Day 0 (Fresh) - IgG | 6703 |
| | CD90+ Day 7 DMSO - IgG | 6278 |
| | CD90+ Day 7 5nM CmpB - IgG | 4005 |

**Supplementary Figure 2: Mitochondrial morphology of CD90⁺ HSCs is altered with MA treatment.**

**A**, Description of chemical structure and primary peptide sequences of mitofusin agonists (MAs) used throughout the study.

**B**, Representative images of resorted $CD90^+$ HSCs stained with Mitotracker Red after 7 days of culture with DMSO, 5nM Cmp A or 5nM Cmp B. Scale bar is 1μm.

**C**, Frequency of mitochondrial lengths in resorted $CD90^+$ HSCs cultured for 7 days with DMSO versus 5nM Cmp A (top) and DMSO or 5 nM Cmp B (bottom); mean of ≥15 fields of cells from n = 3 biological replicates, two-way ANOVA with Bonferroni's multiple comparisons test.

**D**, Frequency of mitochondrial lengths in resorted $CD90^+$ HSCs cultured for 7 days with 1μM Peptide G versus 1μM Peptide S (top) or 1μM Peptide G versus 1μM Peptide D (bottom); mean of ≥15 fields of cells from 3 biological replicates, two-way ANOVA with Bonferroni's multiple comparisons test.

**E**, Representative FACS histogram of intracellular MFN2 expression within freshly isolated $CD90^+$ HSCs or in $CD90^+$ HSCs cultured for 7 days with DMSO or 5nM Cmp B. Identical cells were stained with either anti-MFN2 or IgG control antibodies.

### Supplementary Figure 3

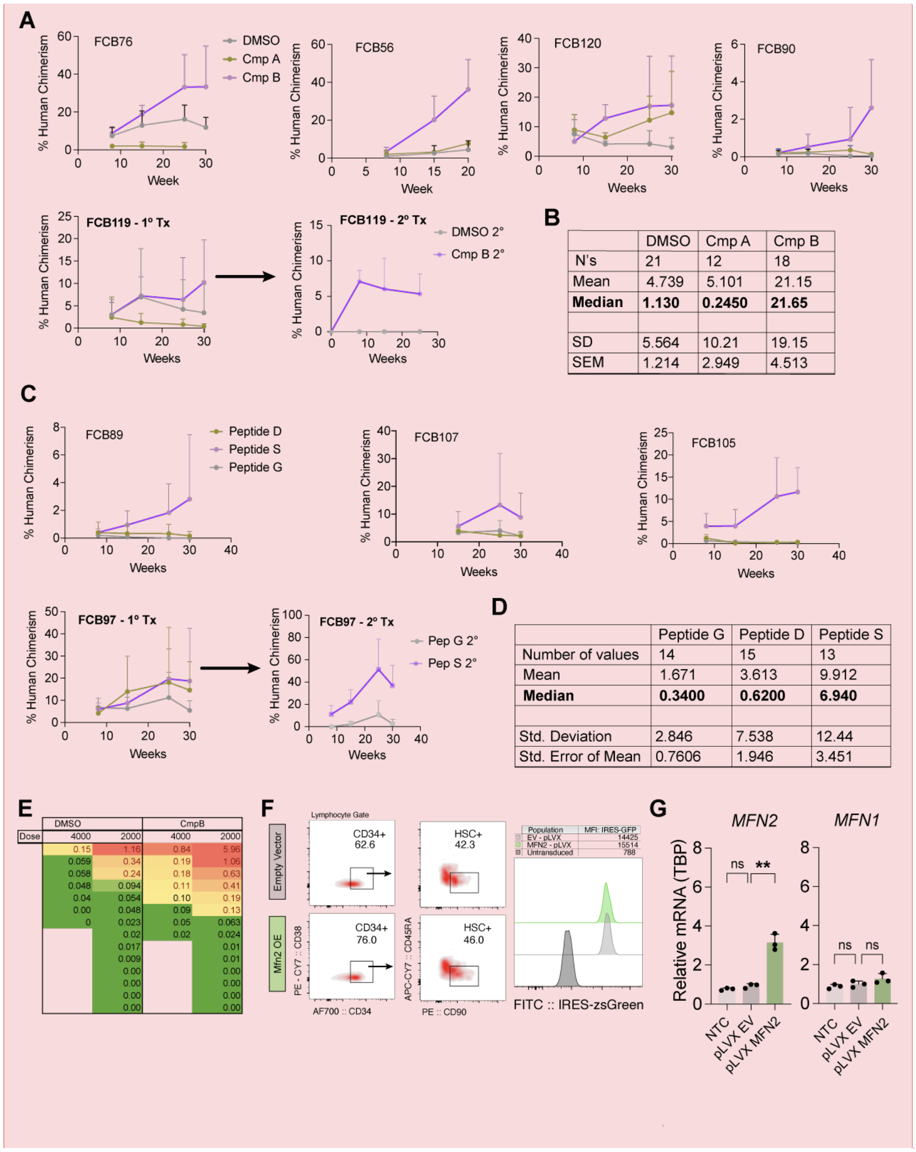

B

| | DMSO | Cmp A | Cmp B |
|---|---|---|---|
| N's | 21 | 12 | 18 |
| Mean | 4.739 | 5.101 | 21.15 |
| **Median** | **1.130** | **0.2450** | **21.65** |
| SD | 5.564 | 10.21 | 19.15 |
| SEM | 1.214 | 2.949 | 4.513 |

D

| | Peptide G | Peptide D | Peptide S |
|---|---|---|---|
| Number of values | 14 | 15 | 13 |
| Mean | 1.671 | 3.613 | 9.912 |
| **Median** | **0.3400** | **0.6200** | **6.940** |
| Std. Deviation | 2.846 | 7.538 | 12.44 |
| Std. Error of Mean | 0.7606 | 1.946 | 3.451 |

E

| | DMSO | | CmpB | |
|---|---|---|---|---|
| Dose | 4000 | 2000 | 4000 | 2000 |
| | 0.15 | 1.16 | 0.84 | 5.96 |
| | 0.059 | 0.34 | 0.19 | 1.06 |
| | 0.058 | 0.24 | 0.18 | 0.63 |
| | 0.048 | 0.094 | 0.11 | 0.41 |
| | 0.04 | 0.054 | 0.10 | 0.19 |
| | 0.00 | 0.048 | 0.09 | 0.13 |
| | 0 | 0.023 | 0.05 | 0.063 |
| | | 0.02 | 0.02 | 0.024 |
| | | 0.017 | | 0.01 |
| | | 0.009 | | 0.01 |
| | | 0.00 | | 0.00 |
| | | 0.00 | | 0.00 |
| | | 0.00 | | 0.00 |
| | | 0.00 | | 0.00 |

**Commented [AB1]:** They want us to change the 3F labels

**Supplementary Figure 3: Long-term engraftment of primary xenografts in NSG recipients.**

**A**, Human chimerism kinetic plots of primary NSG recipients transplanted with 25% of $CD90^+$ HSC cultures treated 7 days with DMSO, 5nM Cmp A, or 5nM Cmp B; n=5 independent experiments. A representative experiment of matched primary and secondary NSG recipient human chimerism kinetics is shown.

**B**, Descriptive statistics of primary NSG recipients 30-weeks post-transplant with 25% of $CD90^+$ HSC cultures treated 7 days with DMSO, 5nM Cmp A, or 5nM Cmp B.

**C**, Human chimerism kinetic plots of primary NSG recipients transplanted with 25% of $CD90^+$ HSC cultures treated 7 days with 1μm Pep G, 1μm Pep D, or 1μm Pep S; n=4 independent experiments. A representative experiment of matched primary and secondary NSG recipient human chimerism kinetics is shown.

**D**, Descriptive statistics of primary NSG recipients 30-weeks post-transplant with 25% of $CD90^+$ HSC cultures treated 7 days with 1μm Pep G, 1μm Pep D, or 1μm Pep S .

**E**, Human chimerism percentages of high cell dose (4,000) and low cell dose (2,000) LDA transplant recipients 15 weeks post-transplant from CBU $CD90^+$ HSCs cultures treated with DMSO or 5nM Cmp B for 7 days.

**F**, FACS phenotype plot of HSC cultures 72 hours after transduction with empty vector or MFN2 lentivirus (left panel). Representative histogram of lentiviral IRES-GFP reporter expression in HSC cultures 72 hours after transduction with empty vector or MFN2 lentivirus compared to no transduction control culture (right panel).

**Commented [AB2]:** We should state transduction efficiency - 100% based on flow.

**G**, Expression of *MFN2* and *MFN1* mRNA in sorted $CD90^+$ HSCs 72 hours after transduction with empty vector, MFN2 lentivirus , or non-transduced control. mRNA levels were quantified by qRT-PCR and normalized to the TBP housekeeping gene; n=3 biological replicates, *P<0.05, one-way ANOVA with Dunnett's post hoc test.

### Supplementary Figure 4

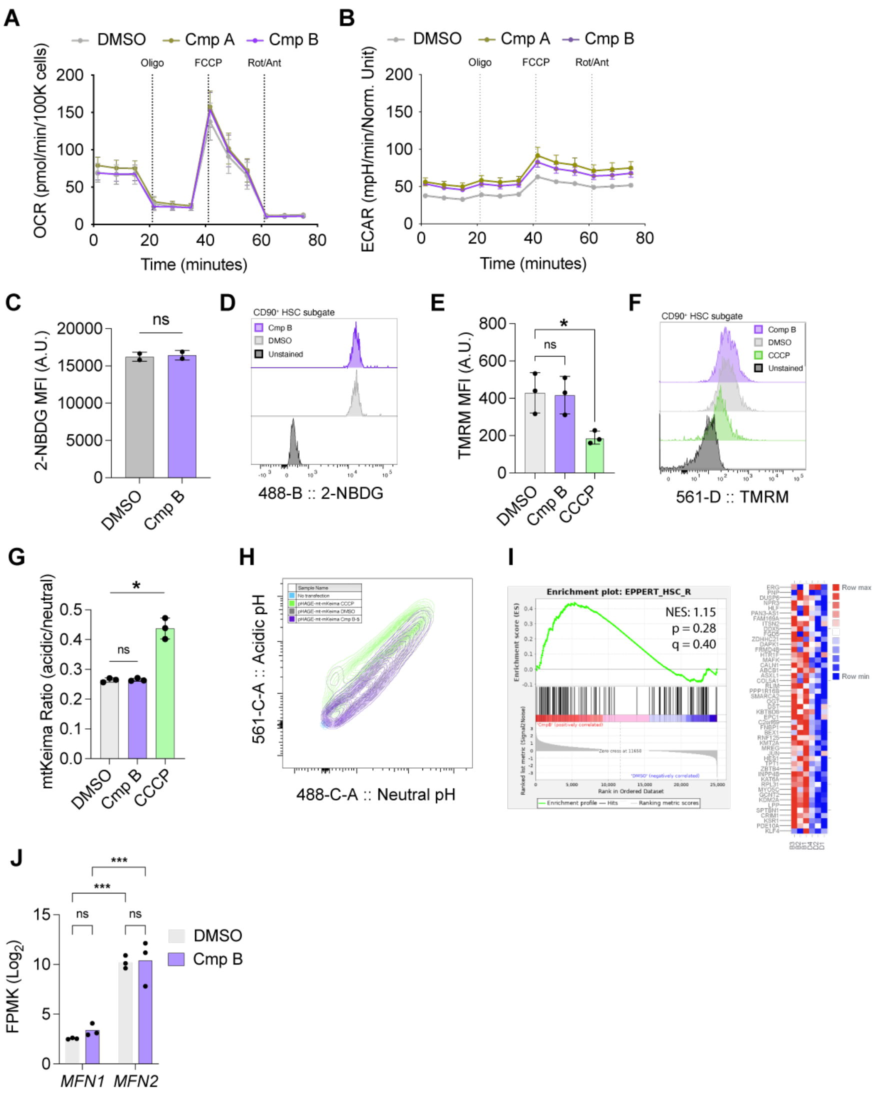

**Supplementary Figure 4: MA treatment does not alter calcium signaling, respiration or mitophagy.**

**A,** Oxygen consumption rate (OCR) derived from Seahorse extracellular flux analysis of cultures initiated with 50,000 $CD34^{+}CD38^{-}CD45RA^{-}$ cells treated for 3 days with DMSO, 5nM Cmp A, or 5nM Cmp B; n=5 independent experiments, one-way ANOVA with Dunnett's post-hoc test.

**B,** Extracellular acidification rate (ECAR) derived from Seahorse extracellular flux analysis of cultures initiated with 50,000 $CD34^{+}CD38^{-}CD45RA^{-}$ cells treated for 3 days with DMSO, 5nM Cmp A, or 5nM Cmp B; n=5 independent experiments, one-way ANOVA with Dunnett's post-hoc test.

**C,** Flow cytometric 2-NBD glucose uptake in phenotypic $CD90^{+}$ HSCs cultured for 3 days with DMSO, 5nM Cmp A, or 5nM Cmp B followed by treatment with 10µM 2-NBDG for 1h; n=5 independent experiments, student's t-test.

**D,** Representative histogram of 2-NBDG signal gated on phenotypic $CD90^{+}$ HSCs cultured for 3 days with DMSO or 5nM Cmp B.

**E,** Flow cytometric TMRM mitochondrial polarization in phenotypic $CD90^{+}$ HSCs cells cultured for 3 days with DMSO, 5nM Cmp B, or treatment with 50µM CCCP 3h prior to experiment followed by loading of 100nM TMRM for 30 minutes; n=3 independent, one-way ANOVA with Dunnett's post-hoc test.

**F,** Representative histogram of TMRM signal gated on phenotypic $CD90^{+}$ HSCs cultured for 3 days with DMSO, 5nM Cmp B, or 50µM CCCP.

**G,** Flow cytometric quantification of mitophagy in HeLa cells transfected with Mito-Keima plasmid cultured for 24 hours with DMSO, 5nM Cmp B, or treatment with 50µM CCCP 3h prior to experiment; n = 3 independent experiments, one-way ANOVA with Dunnett's post-hoc test.

**H**, Representative contour plot of Mito-Keima signal in HeLa cells cultured for 24 hours with DMSO, 5nM Cmp B, or treatment with 50µM CCCP 3h prior to experiment.

**I**, GSEA results of Cmp B versus DMSO RNA-seq dataset against the EPPERT HSC human gene set.

**J,** Expression of *MFN1* and *MFN2* transcripts from Cmp B versus DMSO RNA-seq dataset; *** $p \leq 0.001$, n=3 independent experiments, two-way ANOVA with Tukey's multiple comparison test.

### Supplementary Figure 5

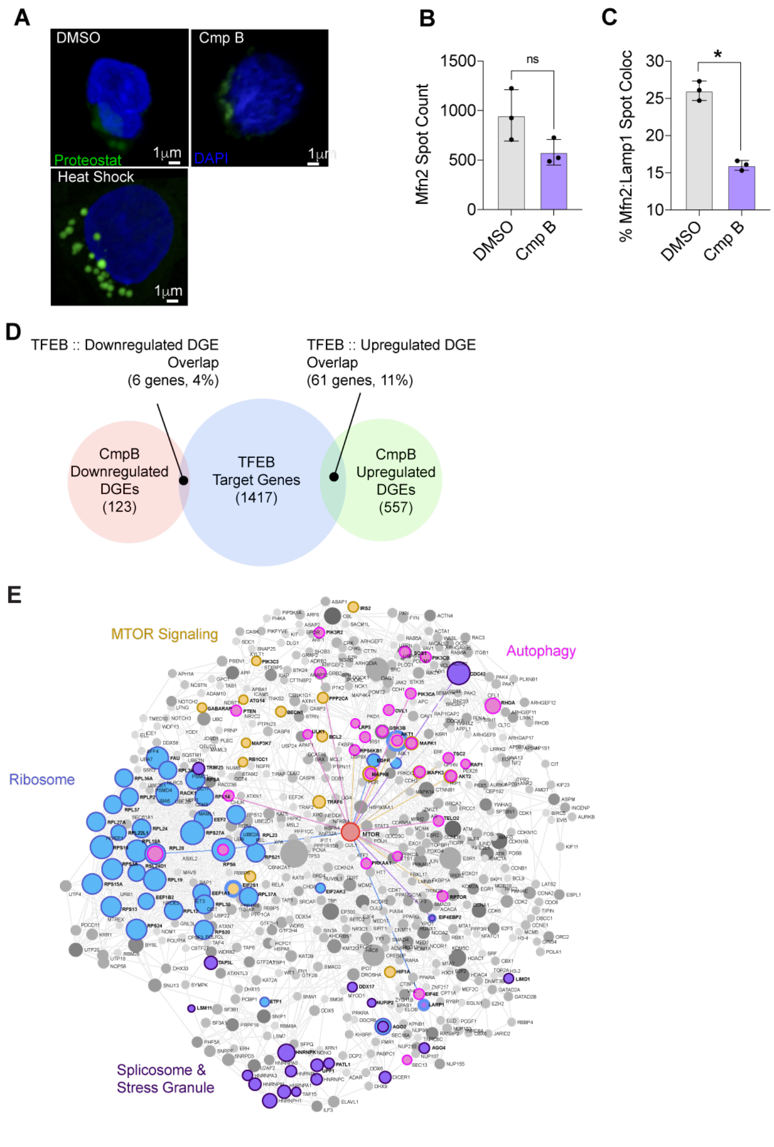

**Supplementary Figure 5: MA treatment upregulates MTOR signal transduction pathways.**

**A**, Representative confocal micrographs of ProteoStat staining in CD90$^+$ HSCs cultured 3 days with DMSO or 5nM Cmp B. Positive control cultures were incubated at 42ºC for 1 hour to induce a heat shock as a positive control. Scale bar is 1μm.

**B,** Quantification of Mfn2 spots in resorted CD90$^+$ HSCs cultured with DMSO or 5nM Cmp B for 3 days. Spot analysis was performed in IMARIS program; n = 3, *p < 0.05, two-tailed student t-test.

**C**, Fraction of Mfn2 spots colocalized with LAMP1 spots in resorted CD90$^+$ HSCs cultured with DMSO or 5nM Cmp B for 3 days. Spot analysis was performed in IMARIS program; n = 3 biological replicates, *p < 0.05, two-tailed student t-test.

**D**, Venn diagram showing overlap between TFEB target genes from the TFEBdb platform database (blue area, 1,417 genes) and RNA-seq differential expressed genes (DEGs) from resorted CD90$^+$ HSC cultured for 7 days with DMSO or 5nM Cmp B. Total downregulated DEGs (red area, 123 genes) and total upregulated (green area, 557 genes) are shown. Overlapping areas shown hits between Cmp B DEG lists and TFEB targets.

**E**, STRING Interactome network analysis of generic protein-protein interactions using upregulated DEGs in resorted CD90$^+$ HSCs cultured for 7 days with DMSO or 5nM Cmp B. A confidence score threshold of 920 was used requiring experimental evidence.

### Supplementary Figure 6

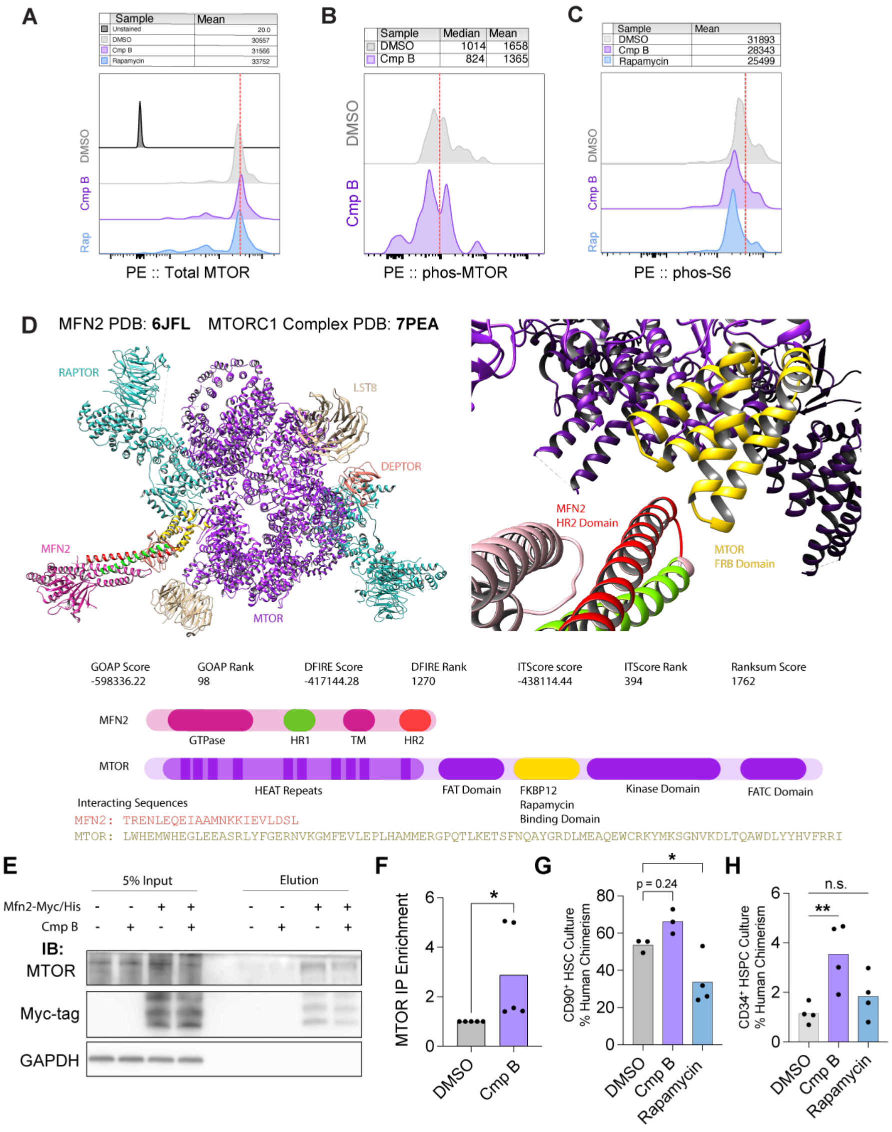

**Supplementary Figure 6: Putative protein interaction with MFN2 correlates with inhibition of MTOR signaling.**

**A,** Representative histogram of total mTOR signal gated on phenotypic $CD90^+$ HSC cultures after culture for 3 days with DMSO or 5nM Cmp B.

**B,** Representative histogram of phospho-mTOR signal gated on phenotypic $CD90^+$ HSCs after culture for 3 days with DMSO or 5nM Cmp B.

**C,** Representative histogram of phospho-S6 signal gated on phenotypic $CD90^+$ HSCs after culture for 3 days with DMSO or 5nM Cmp B.

**D**, LZerD protein-protein docking simulation results using Mfn2 (PDB: 6JFL) and MTORC1 complex (PDB: 7PEA). HR1 and HR2 domains of MFN2 (red and green) and FBD domain of MTOR (yellow) depict interaction between the two proteins. Ranksum score of GOAP, DFIRE and ITScore ranks and schematic map of MFN2 and MTOR protein domains are shown.

**E**, Representative western blot of MTOR immunoprecipitation in 293T cells transfected with MFN2-MYC/HIS plasmid followed by 24h treatment with DMSO or 5nM Cmp B.

**F**, Densitometry analysis of mTOR immunoprecipitation levels in 293T cell lysates transfected with MFN2-MYC/HIS plasmid followed by 24h treatment with DMSO or 5nM Cmp B. Data was normalized to the DMSO control; n = 5, *$p$ > 0.05, student's *t*-test.

**G,** Human chimerism of NSG recipients transplanted with 25% of $CD90^+$ HSC cultures treated for 7 days with DMSO, 5nM Cmp B, or 200nM rapamycin 15 weeks post-transplant; n=3-4 mice, * p<0.05; one-way ANOVA with Dunnett's post hoc test.

**H,** Human chimerism 8 weeks post-transplant from NSG recipients transplanted with 25% of cultures initiated with 60K phenotypic $CD34^+$ cells cultured for 7 days with DMSO, 5nM Cmp B, or 200nM rapamycin; n=4 mice. (* p<0.05; one-way ANOVA with Dunnett's post hoc test).

**Supplementary Figure 7**

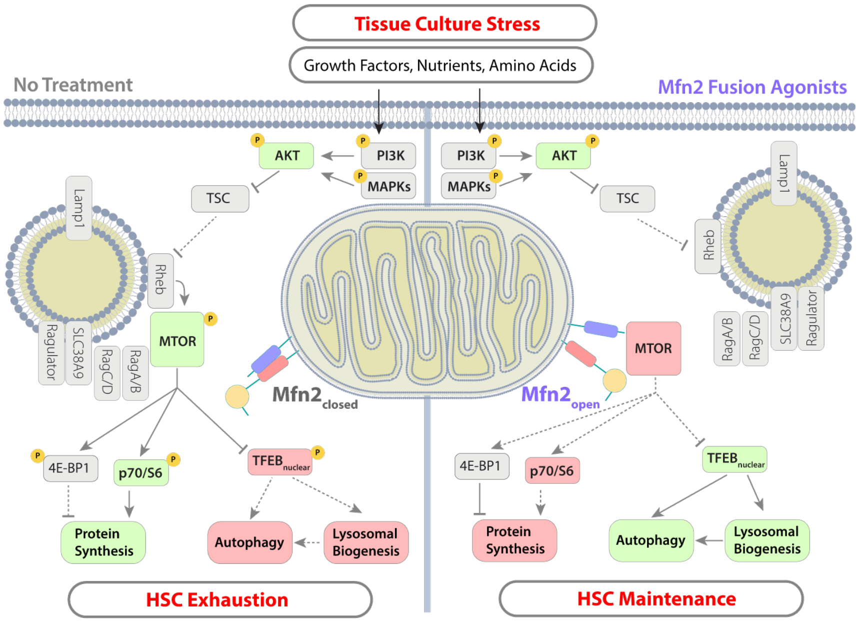

**Supplementary Figure 7: Schematic diagram of between MFN2 and MTOR in the presence of MA treatment.**

In the presence of tissue culture stress (high levels of growth factors, nutrients, amino acids, etc.) upstream mitogenic signaling activates AKT leading to downstream anabolic programs. Activation of MTOR leads to enhanced protein synthesis and reduced autophagic flux and correlates with exhaustion of HSCs *in vitro*. In the presence of MAs, fusion competent MFN2 binds the FRB domain and sequesters MTOR from the activation site on lysosomes. Consequentially, protein translation is reduced, and lysosomal biogenesis and autophagy is activated thus preserving HSC potency during exposure for MAs during *in vitro* expansion.

**Supplementary Table 1: Differentially expressed genes in Cmp B-treated CD90$^+$ HSCs.**

RNA-seq differential expressed genes (DEGs) lists from resorted CD90$^+$ HSC cultured with DMSO or Cmp B for 7 days. List cutoff $p \leq 0.05$.

**Supplementary Table 2: DEG pathway analysis in Cmp B-treated CD90$^+$ HSCs.**

KEGG pathway and Gene Ontology (GO) term analysis of RNA-seq differential expressed genes (DEGs) lists from resorted CD90$^+$ HSC cultured with DMSO or Cmp B for 7 days .

**Supplementary Table 3: TFEB target gene list correlation to DEGs from Cmp B-treated CD90$^+$ HSCs**

List of genes overlapping between TFEB target genes and RNA-seq differential expressed genes (DEGs) lists from resorted CD90$^+$ HSC cultured with DMSO or Cmp B for 7 days .

**Supplementary Methods**

*Cord Blood Cell Isolation*

CBUs were thawed at 37ºC until ice crystals were absent. Volume-reduced CBUs (30mL) were flushed with a needle and syringe and washed in 600mL of 1x PBS and spun at 850xg at 4ºC for 25min. The supernatant and RBC layers were aspirated, and cell pellets pooled for an additional washing with 50mL of PBS. After washing, cells were counted and incubated with CD34 microbeads (Miltenyi Biotec) according to manufacturer's protocol with the addition of 50kUz of DNase I (Thermo Fisher). Cells were incubated with rotation at 4ºC for 30min followed by washing with PBS. Labeled CD34+ cells were eluted using the AutoMACS POSSELD2 program (Miltenyi Biotec), counted and stained with the human HSPC antibody cocktail. Phenotypic HSCs ($Lineage^-$, $CD45^{low}$, $CD34^+$, $CD38^-$, $CD45RA^-$, $CD90^+$) were then sorted using a BD FACSAria Fusion cell sorter and used for experiments.

*Peripheral Blood and Bone Marrow Isolation*

To isolate peripheral blood mononuclear cells (PBMCs), mice were bled submandibular and whole blood was treated twice with 1x Ammonium-Chloride-Potassium (ACK) Lysing Buffer to lyse RBCs. PMBCs were stained with a peripheral blood antibody cocktail (*see antibodies table*) for 20 minutes on ice and washed cells were immediately analyzed by flow cytometry. To isolate BM cells, recipient femurs and tibias were pooled and crushed using a sterile pestle in PBS. Cell supernatants were passed through a 40μm filtration followed the ACK lysis. Cells stained with a human HSPC antibody cocktail (*see antibodies table*) for 20min on ice, washed and analyzed by flow cytometry. FCS Express 7 Research and FlowJo 10 software were used to analyze all data from flow cytometry.

*Hematopoietic Stem Cell Culture*

Human HSCs cultures were carried out in complete medium using StemSpan SFEM (STEMCELL Technologies) as the basal media and reconstituted with addition of Penicillin-Streptomycin (1X), GlutaMAX™ Supplement (1X), hrTPO 100ng/mL, hrSCF 100ng/mL, hrFLT-3 100ng/mL, hrIL-3 50ng/mL and hrIL-6 20ng/mL. $CD90^+$ HSC cultures were initiated with 10,000 phenotypic $CD90^+$ HSCs in 96 U-bottom well plates (5,000 cells per well). $CD34^+$ HSC cultures were initiated with 60,000 phenotypic $CD34^+$ HSPCs in 12-well well plates. To test small molecule MA function, complete media was supplemented with 5nM Compound A, 5nM Compound B or DMSO as control (treatment <0.01% v/v). To test peptide MA function, complete media was supplemented with 1μM Peptide-S, Peptide-D, or 1μM Peptide G as control (treatment <0.01% v/v). Cells were cultured for 7 days in 5%$CO_2$ and 5%$O_2$ at 37°C. Half of the media volume was exchanged every other day. At the indicated timepoint, culture cell counts from each culture condition was obtained, as well as a FACS phenotypic human HSC flow analysis to determine phenotypic HSC count yields.

*Cell Line Maintenance and MFN2 KO Cell Line Generation*

293FT and HeLa cells (ATCC) were sub-cultured in 10% FBS/DMEM with 1x pen/strep in a 5% $CO_2$ incubator at 37°C. Engineering of MFN2 KO cell lines was designed using the TrueDesign Genome Editor software according to manufacturer's instructions

(ThermoFisher). 50,000 293T cells were seeded 10%FBS/DMEM and 7.5pmol of Cas9 protein (Invitrogen Cat. No. A36497), 7.5pmol MFN2 sgRNA, and 500ng of each TruTag donor DNA (Invitrogen Cat. No. A53815) were delivered via transfection using Lipofectamine CRISPRMAX reagent and Lipofectamine Cas9 Plus reagent overnight. After 72h, transfected cells were sorted for expression of both RFP and GFP indicating incorporation of guide edits into both MFN2 genomic alleles. To eliminate GFP/RFP reporters, double positive RFP/GFP transfected cells were treated with 500ng Cre recombinase mRNA (Trilink Cat. No. L-7211) using Lipofectamine CRISPRMAX as instructed by the manufacturer. Decay of fluorescence signal was monitored by fluorescence microscopy and flow cytometry over 7 days. Double-negative GFP/RFP cells, indicating removal of the fluorescent and selection cassette, were then used for all experiments. Validation of KO in the GFP/RFP double negative cells was confirmed by western blotting for MFN2.

*Lentivirus Production and HSPC Transduction*

Lentiviral particles were produced by seeding 293T cells at $7\times10^5/cm^2$ in DMEM serum-free media (Lonza, Basel, Switzerland) overnight followed by PEI mediated transfection of viral packaging vectors (pMD2.G and pPAX2) and expression construct in the pLVX-EF1$\alpha$-IRES-zsGreen backbone vector (CloneTech). Media were pooled after 36–48h, clarified and concentrated by ultracentrifugation (100,000xg), resuspended in DMEM media and stored at −80°C. Virus titer was calculated from transduction of NIH-3T3 fibroblasts with serial dilutions of the viral. Sorted $CD90^+$ HSCs were transduced with either empty vector (EV-pLVX) or MFN2-pLVX lentivirus at a MOI of ≥150 in the presence of 6µg/mL polybrene (Sigma) and spun at 900xg for 20min at 20°C. Supernatant was aspirated and replaced with complete media and cultured overnight. Transduction efficiency was assessed 48-72h post-transduction by measuring IRES-GFP expression via flow cytometry.

*HSPC Autophagic Flux Assay*

CD34+ HSPCs were transfected with FUW mCherry-GFP-LC3 plasmid (Addgene 110060) by electroporation performed using the Lonza 4D-Nucleofector system with the P3 Primary Cell 4D-Nucleofector Kit according to the manufacturer's instructions. Cells were electroporated with 1 µg plasmid per 1 × $10^6$ cells using the $CD34^+$ cell electroporation program, cultured overnight, and then treated with DMSO, 5 nM Cmp B or 200nM rapamycin for the indicated experimental duration. Cells were mounted to CellTak treated glass bottom wells, fixed with 4% PFA/PHEM buffer and mounted with Vectashield mounting medium. SCRM images of transduced cells were collected on a Leica DM8000 deconvolution microscope at 63x magnification. The ratio of mCherry to GFP ratio for individual organelle structures were collected and the mean ratio per cell was calculated.

*TFEB Luciferase Gene Reporter Assay*

CD34+ HSPCs were transfected with equal amounts of TFEB luciferase reporter pLminP-Luc2P-RE28 plasmid (Addgene 90370) and pSV-βGal (Promega E1081) by electroporation and was performed using the Lonza 4D-Nucleofector system with the P3 Primary Cell 4D-Nucleofector Kit according to the manufacturer's instructions. Cells were electroporated with 1 μg plasmid per 1 × $10^6$ cells using the $CD34^+$ cell electroporation program, cultured overnight, and then treated with 5 nM Cmp B or DMSO for the indicated experimental duration. Cells were lysed in 100uL of 1x Reporter Lysis Buffer (Promega) and subject to freeze/thaw lysis at -80C. Lysates were incubated with 100uL of Luciferase Assay Buffer (Promega E1501) using a Cytation 5 plate reader (Biotek). Normalized β-gal activity was measured using the BetaGlo kit (Promega E4720) according to manufacturer's instructions. The ratio of Luciferase to β-Gal expression was calculated to report relative gene expression activity.

*Mitophagy Measurements in HeLa cells*

For mitophagy experiments, 50,000 HeLa cells were plated in 12-well dishes and transfected with pHAGE-Mito-mKeima (Addgene 131626) overnight . The next day, media was changed to 1% fetal bovine serum/DMEM and treated with for 24h with DMSO, 5nM Cmp B, or treated with 50μM CCCP 3h prior to experiment as positive control. Cells were trypsinized and analyzed by flow cytometry to calculate the ratio of neutral-to-acidified mitochondria.

*TFEB-GFP nuclear localization experiments in HeLa cells*

For TFEB nuclear localization experiments, 50,000 HeLa cells were plated in 6-well glass-bottom dishes (Cellvis) and transfected with pEGFP-N1-TFEB plasmid overnight. Cells were cultured for 24h with DMSO, 5nM Cmp B, 200nM rapamycin or treated with 50μM CCCP 3h prior to experiment as positive control. Cells were fixed with 4% PFA/1xPHEM buffer for 15min. Cells were mounted and nuclear counterstained with Prolong Diamond

Antifade DAPI (Thermo Fischer). Fluorescent images were acquired with a Leica microscope using a 40x objective and the percentage of TFEB-GFP localized to the nucleus was calculated.

*Western Blots*

For total cell lysate experiments, 293T or HeLa cultures were lysed in RIPA buffer, 50mM Tris pH 7.5, 137mM NaCl, 0.1% SDS, 0.5% deoxycholate and protease inhibitors (Roche). Lysates were incubated on ice for 10 minutes and sonicated for 30 seconds 50% power. Lysates were centrifuged at 15,000xg for 10 minutes and supernatant was stored at -80C. All protein samples were denatured in 4x sample buffer at 95 °C and loaded onto 4–12% Bis-Tris SDS-PAGE gradient gels (Invitrogen). Gels were transferred onto 0.22μm nitrocellulose membrane and stained with Ruby Red (Molecular Probes, Carlsbad, CA) to confirm transfer. Membranes were blocked with BSA in 0.3%Tween-20/TBS and incubated with anti-MFN2 (1:200), anti-mTOR (1:500), anti-Phos-S6 (1:200), anti-Phos-AKT (1:500), anti-GAPDH (1:1000), anti-His tag (1:500), and anti-β-actin (1:5000) overnight. Membranes were washed, incubated with HRPO-conjugated secondary antibodies for 1 hour. After incubation, membranes were developed with Super Signal West Fempto ECL reagent (Pierce).

*Immunoprecipitation Assays*

For endogenous MFN2 co-immunoprecipitation, 293FT cells were plated to ~70% confluency on 100-mm culture dishes for 24 hours. Cells were harvested by scraping in Pierce lysis buffer supplemented with HALT protease and phosphatase inhibitor cocktail, and lysates were clarified by centrifugation at 600 × g for 3 minutes. Endogenous MFN2 was immunoprecipitated from clarified lysates using a mouse anti-MFN2 antibody (Abcam ab56889), with species-matched IgG used as a negative control. Immune complexes were captured using Protein A/G beads, washed, eluted, and analyzed by immunoblotting with a rabbit anti-MFN2 antibody (Cell Signaling Technologies 9482S).

For epitope-tagged precipitation, HeLa cells were plated to 70% confluency on 100 mm culturing dishes and transfected with 6x-his tagged MFN2 plasmid using Lipofectamine 3000 (Thermo Scientific) for 48 hours. Cells were cultured with either Cmp B or DMSO control for 24 hours and then harvested with Peirce Lysis buffer (Promega) supplemented with HALT™ protease phosphatase inhibitor cocktail (Thermo Scientific) through scrapping. Cells were centrifuged at 600 x g for 3 minutes. The supernatant was recovered and applied to HisPur™ cobalt resin columns for 1 hour with rotation at 4˚C. Immunoprecipitation was performed using Peirce™ His protein interaction pulldown kit (Thermo Scientific) as described by the manufacturer. Eluted complexes were analyzed by immunoblotting for epitope MFN2 with a mouse anti-Myc antibody (Sigma 05-724).

*Antibody Immunofluorescence*

Freshly isolated CBU populations or cultured phenotypic HSCs as described were sorted in complete media and plated into glass-bottom 96-Well plates (Cellvis) coated with CellTak (Corning, 30mg/cm$^2$). Cells were adhered for 30 min at RT and fixed with 4% PFA/2xPHEM buffer for 15min. Cells were then permeabilized with 0.1% Triton X-100/PBS for 5min and blocked with 1% BSA/PBS for 1 hour. Cells were incubated with primary antibody overnight, washed and incubated with secondary antibodies for 2 hours the next day. Cell nuclei were counterstained and mounted with Prolong Diamond Antifade DAPI (Thermo Fisher). Images were acquired with a Zeiss LSM880 microscope using a 100x objective and analyzed using Imaris 11.1.

*Mitotracker Red (MTR) Mitochondrial Length Measurements*

CD90$^+$ HSCs (10,000 cells per condition) were cultured for 7 days in DMSO or with 5nM Cmp A or 5nM Cmp B. Phenotypic CD90$^+$ HSCs were re-sorted and incubated in complete medium containing 250nM MitoTracker Red (Thermo Fisher Scientific) at 37°C for 30 minutes. Cells were adhered to CellTak-coated 96-well glass-bottom plates, fixed in 4% paraformaldehyde prepared in 2× PHEM buffer, and counterstained/mounted with ProLong Diamond Antifade mounting medium with DAPI. Images were acquired using a Zeiss LSM880 confocal microscope with a 63× objective. Mitochondrial lengths were measured with Fiji or Imaris 11.0.

*Image Quantification*

For mitochondrial length measurements, confocal or deconvoluted z-stacks were collected and projected as a z-project in ImageJ (NIH, Bethesda, MD). Individual mitochondria were manually traced, binned into length categories, and expressed as percent of cellular mitochondria. The mean ±SEM number of mitochondria falling into each length category collected from ≥15 fields (30-50 cells) are expressed. For NFAT nuclear localization quantification, confocal z-stacks were collected and a 0.5mm in the center of the cell was projected as a z-project in ImageJ. Nuclear boundaries were constructed using DAPI staining. The ratio of staining within the nuclear boundary to total staining was expressed as percent of NFAT signal. The mean ±SEM for ≥10 fields (20-40 cells) are expressed. For immunofluorescence intensity measurements, confocal or deconvoluted z-stacks were collected and projected as a z-project in ImageJ. Thresholds were set based on IgG-stained negative control cells and the integrated density value of each signal per cell was recorded. The mean ±SEM for ≥15 fields (30-50 cells) are expressed.

*Flow Cytometry Dye Quantifications*

To measure glucose uptake, cells were treated with 200 μM 2-NBDG at 37ºC for 2 hours. Cells were washed with PBS and stained with surface antigens human antibodies and analyzed on flow cytometer with 488-nm excitation. To measure lysosome quantity and acidification, cells were cultured under conditions as described and then treated with 100nM Lysotracker Green (Thermo Fisher) at 37ºC for 45min and 30min, respectively. As a control, cells were treated with 200nM rapamycin or 50μM chloroquine 18 hours before the experiment. Cells were then washed with PBS and stained with surface marker antibodies and analyzed by flow cytometry. To evaluate mitochondrial membrane potential, cells were cultured under conditions as described and then treated with 20nM MitoProbe TMRM reagent (Thermo Fisher) for 30 min (incubator 37ºC, 5% O2 and 5% CO2). Cells were washed and stained with surface marker antibodies and analyzed by flow cytometry. To measure protein synthesis rates, Click-iT® OP-puromycin reagent (Thermo Fisher) was used according to manufacturer's instructions. Briefly, cells were cultured under conditions as described followed by treatment, stained with surface marker antibodies, and then fixed and permeabilized using 4% PFA/PHEM and 0.1% Triton X-

100/PBS. Cells were then incubated with the 1x Detection Cocktail (Alexa-Fluor-488 conjugated to OPP) for 30 min at room temperature, washed and analyzed by flow cytometry. To measure protein aggresome formation, the PROTEOSTAT Protein aggregation assay (Enzo Life Sciences) was used according to manufacturer's instructions. As a positive control, cells were incubated at 42ºC for 3 hours before the experiment. After culture, cells were incubated for 20 min in an antibody cocktail, washed with PBS, fixed and permeabilized using 4% PFA/PHEM and 0.1% Triton-X 100/PBS and then incubated with 1x Aggresome Detection Reagent for 30 min at room temperature. Results were analyzed by flow cytometry. FCS Express 7 Research or FlowJo software were used to analyze flow cytometry data.

*Seahorse Metabolic Flux Experiments*

For all Seahorse metabolic assays, DMEM without sodium bicarbonate or HEPES buffer and neutralized to pH 7.4 at 37ºC (Buffer-free DMEM) immediately prior to the experiments. XFp flux cartridges were hydrated in XF Calibrant overnight. For all assays, 50,000 HSPCs ($CD34^+CD38^-CD45RA^-$ cells) were cultured with small molecule MAs or DMSO control for 3 days and the entire culture was then plated onto one well of a Seahorse XFp culture plate coated overnight with Cell-Tak reagent. Cells were immobilized by centrifugation at 20xg for 5min at RT and washed twice with DMEM media. Cells were equilibrated in fresh bicarbonate-free DMEM in a humidified non-CO2 incubator and assayed within 30min to 1 hour after cell culture harvest. Flux cartridges were loaded with respiration inhibitors prepared with Buffer-free DMEM according to manufacturer's instructions. Oxidative phosphorylation and glycolysis values were obtained using the XFp Mito Stress Kit and Glycolysis Stress Kit, respectively. Metabolic parameters were derived from calculations based on manufacturer's instructions. Due to feasibility of cell number isolation, experiments are represented as one technical replicate per cell type, per condition over 4 independent experiments.

Gene Set Enrichment Analysis (GSEA)

GSEA was performed using the fgsea R package and the fgseaMultilevel() function. The log2 fold change from the DMSO vs Cmp B differential expression comparison was used to rank genes. Gene set collections from the Molecular Signatures Database (MSigDB) were curated using the msigdbr R package. Prior to running GSEA, the list of gene sets was filtered to include only gene sets with between 5 and 1000 genes. GSEA results for each tested gene set are shown in the table. The Adj p-value column contains the false discovery rate (FDR)-adjusted p-value. The NES column contains the normalized enrichment score (NES) computed by GSEA, which represents the magnitude of enrichment as well as the direction. Enrichment plots show the genes ranked by log2 fold change along the x-axis with the vertical ticks representing the location of the genes in the gene set. The heatmap displays the expression of the genes, red being more expressed in the first group (High) and blue more expressed in the second group (Low). The green line shows the enrichment score. Analyses performed using Pluto (https://pluto.bio).

*Network Analyst of RNA-seq DGE lists*

The Network Analyst web server was used to perform systems interpretation of RNA-seq differential gene expression (DGE) data through knowledge-based protein-protein interaction (PPI) networks. Briefly, statistically significant RNA-seq DGE lists were analyzed for generic PPI using the STRING Interactome with the requirement for experimental evidence with a confidence score cutoff of 920. The minimum network option was utilized to visualize the network such that seed proteins and essential non-seed proteins were kept to maintain the network connectivity, simplify network density and identify key connectivity. (https://www.networkanalyst.ca/NetworkAnalyst)

*Protein Docking*

The Local 3D Zernike descriptor-based Docking algorithm (LZerD) pairwise docking program was used to predict protein-protein interactions between MFN2 and MTOR. Using geometric hashing to generate ligand orientations, LZerD utilizes 3D Zernike descriptors as shape matching criteria. Scoring is based on a shape complementarity term defined by the local shape Zernike and orientation of surfaces. Crystal structures for human MFN2 (PDB: 6JFK) and the human MTORC1 complex (PDB: 7PEA) were used without constraints and the top scoring model was selected for interpretation.
